# Glutamate signaling and neuroligin/neurexin adhesion play opposing roles that are mediated by major histocompatibility complex I molecules in cortical synapse formation

**DOI:** 10.1101/2024.03.05.583626

**Authors:** Gabrielle L. Sell, Stephanie L. Barrow, A. Kimberley McAllister

## Abstract

Although neurons release neurotransmitter before contact, the role for this release in synapse formation remains unclear. Cortical synapses do not require synaptic vesicle release for formation ^1-4^, yet glutamate clearly regulates glutamate receptor trafficking ^5,6^ and induces spine formation ^7-11^. Using a culture system to dissect molecular mechanisms, we found that glutamate rapidly decreases synapse density specifically in young cortical neurons in a local and calcium-dependent manner through decreasing NMDAR transport and surface expression as well as co-transport with neuroligin (NL1). Adhesion between NL1 and neurexin 1 protects against this glutamate-induced synapse loss. Major histocompatibility I (MHCI) molecules are required for the effects of glutamate in causing synapse loss through negatively regulating NL1 levels. Thus, like acetylcholine at the NMJ, glutamate acts as a dispersal signal for NMDARs and causes rapid synapse loss unless opposed by NL1-mediated trans-synaptic adhesion. Together, glutamate, MHCI and NL1 mediate a novel form of homeostatic plasticity in young neurons that induces rapid changes in NMDARs to regulate when and where nascent glutamatergic synapses are formed.

## Introduction

Formation of synaptic connections is an intricate process that is integral to the establishment of the complex circuitry underlying perception, cognition, and behaviour. Impairment of this process is associated with neurodevelopmental and psychiatric disorders ^12-17^. During glutamatergic synapse formation, pre- and postsynaptic proteins accumulate within minutes of stabilized axo-dendritic contact ^14,18-21^. Recruitment of transport packets containing ionotropic glutamate receptors to nascent synapses is integral to this process. N-methyl-D-aspartate receptor (<u>N</u>MDA<u>R</u>) transport <u>p</u>acket<u>s</u> (NRTPs) rapidly deliver NMDARs to newly forming synapses following initial axo-dendritic contact ^21^ via a neuroligin 1 (NL1) -dependent mechanism ^22,23^ and this recruitment is developmentally regulated ^24-26^. NRTP transport along dendrites occurs rapidly and bi-directionally, interspersed frequently with pauses at sites where NMDARs cycle with the plasma membrane and where the initial complement of synapses selectively form ^27^. Periodic delivery to the dendritic surface raises the intriguing possibility that NMDARs are capable of sensing glutamate during their transport before synaptic contacts are established. Yet, the consequences of NMDAR activation for receptor trafficking and synapse formation remain unknown.

During the period of synaptogenesis, axonal growth cones are capable of releasing neurotransmitter before they contact other cells ^18,28-32^. In cortical neurons, vesicles containing the glutamate transporter vGlut1, rapidly cycle along the axonal membrane and at the axonal growth cone ^30,32,33^. However, in contrast to the unambiguous role for neural activity in refinement of connections during critical periods ^34,35^, the role for neurotransmitter release in the initial formation of synaptic connections is unclear. Decades of research indicate that synapse formation is independent of activity because synapses can form in the absence of neurotransmitter release *in vitro* and *in vivo* ^1-4,36^. Yet, glutamate clearly induces dendritic filopodial extension toward axons in culture as well as dendritic spine formation in intact tissue ^7-9^, especially during early stages of synapse formation ^10,11^. Moreover, uncaging of glutamate onto dendrites in cortical slices directly induces the *de novo* formation of dendritic spines with synapses in an NMDAR- and NL-1-dependent manner ^10,37^ and GABA uncaging is sufficient for inducing both dendritic spine formation and clustering of postsynaptic inhibitory proteins ^11^.

This proposed positive role for glutamate in inducing synapse formation at cortical synapses is surprising for several reasons. First, neurotransmitter plays the opposite role at the neuromuscular junction (NMJ). At the NMJ, acetylcholine (ACh) causes internalization of ACh receptors (AChRs) and is opposed by an anti-dispersal signal during NMJ formation ^38,39^. Moreover, glutamate activation of NMDARs at the NMJ during the initial stages of synaptogenesis causes synapse pruning ^40^. Second, even in the central nervous system (CNS), there is substantial evidence that activation of NMDARs causes their internalization at mature synapses ^5,41,42^ and NMDARs in young neurons undergo substantial constitutive and glutamate-induced internalization ^6^. Finally, our lab has shown that a global reduction in activity over days dramatically increases synapse density exclusively in young cortical neurons. Importantly, this homeostatic response requires activity-dependent modulation of major histocompatibility complex I (MHCI) molecules ^43^, a classical immune molecule family found on neurons ^44^ that negatively regulate glutamatergic synaptic density and function^43,45^.

Here, using a reduced cell culture approach essential for examining sufficiency and molecular mechanisms, together with a combination of time-lapse imaging, electrophysiology, immunocytochemistry, confocal microscopy, and biochemical methods, we tested the hypotheses that glutamate at CNS synapses may act in a similar way as acetylcholine at the NMJ—dispersing NMDA receptors and causing synapse elimination unless opposed by trans-synaptic adhesion—and that MHCI molecules mediate this process. We found that acute pharmacological activation of NMDARs and stimulation with glutamate, globally and locally, decrease the mobility and surface expression of NMDARs, the proportion of mobile NMDARs that are transported with NL1, and the density of synapses. NMDAR blockade has the reverse effect. The glutamate-induced decrease in NMDAR transport is local and dependent on NMDAR-mediated Ca^2+^ influx. Importantly, these effects of glutamate are prevented specifically at sites of association of the NMDAR/NL1 complex with Nrxn1 in a hemi-synapse assay. Finally, we determined that major histocompatibility complex I (MHCI) is required for the glutamate-induced synapse loss, through a novel negative regulation of NL1 protein levels.

Together these data show that young cortical neurons undergo a new type of homeostatic plasticity that involves surprisingly rapid changes in glutamatergic synapse density, as opposed to the activity-induced changes in synaptic strength that occur in older neurons ^46^. This new form of plasticity is mediated by MHCI molecules on neurons. Moreover, glutamate release appears to regulate synapse formation by destabilizing postsynaptic components (NMDAR/NL1) that fail to make contact with presynaptic partners (Nrxn1), similar to the role for glutamate in synapse loss ^40^ as well as the opposing roles for acetylcholine (ACh) and agrin ^38^ in synapse formation at the neuromuscular junction (NMJ).

## STAR METHODS

### Key resources table

| REAGENT or RESOURCE | SOURCE | IDENTIFIER |
| --- | --- | --- |
| **Antibodies** |  |  |
| Mouse anti-vGlut1 (1:1000) | Chemicon | AB5905 |
| Rabbit anti-GluN2A (1:400) | Upstate<br>Biotechnology | 07-632 |
| Rabbit anti-GluN2B (1:400) | Chemicon | AB1557 |
| Rabbit anti-GFP (1:1000) | Molecular Probes | A11122 |
| Goat anti-mCherry (1:2000) | Sicgen | AB0040-200 |
| Rat anti-HA (1:1000) | Roche | Clone 3F10 |
| Rabbit anti-VAMP (1:1000) | Synaptic Systems | 104-202 |
| Mouse anti-PSD95 (1:750) | Antibodies, Inc. | K28/43 |
| Donkey anti-rabbit 680 | LI-COR | 926-68073 |
| Donkey anti-mouse 800 | LI-COR | 926-32212 |
| Donkey anti-goat 800 | LI-COR | 926-32214 |
| AlexaFluor-488 anti-rabbit (1:400) | Invitrogen | A-21206 |
| AlexaFluor-405 anti-mouse (1:400) | Invitrogen | A-31553 |
| Cy3-conjugated donkey anti-goat (1:400) | Jackson Imm. | 705-165-147 |
| **Chemicals, peptides, and recombinant proteins** |  |  |
| N-methyl-D-aspartic acid (NMDA) | Sigma | M3262 |
| glutamate (L-glutamic acid) | Sigma | G8415 |
| D(-)-2- amino-5 phosphopentanoic acid (APV) | Sigma | A8054 |
| 6-cyano-7-nitroquinoxaline-2,3-dione (CNQX) | Sigma | C127 |
| α-methyl-4-carboxyphenylglycine (MCPG) | Sigma | M4796 |
| TTX | Abcam | ab120055 |
| Poly-L-lysine | Sigma | P2636 |
| Mycozap | Lonza | LT07-818 |
| Fetal Bovine Serum | Gemini | 100-106 |
| B27 plus | ThermoFisher | A3582801 |
| Glutamax | ThermoFisher | 35050061 |
| Sodium Pyruvate | ThermoFisher | 11360070 |
| HEPES | ThermoFisher | 15630080 |
| Lipofectamine 2000 | Invitrogen | 11668019 |
| **Experimental models: Organisms/strains** |  |  |
| Neurologin-1 knockout mouse model | Craig Powell | 47 |
| H2-Kb/H2-Db double knockout mouse model | Lisa Boulanger | 48,49 |
| β2m knockout mouse model | Jackson Labs | B6.129P2-<br>β2m<sup>tm1Unc</sup>/J |
| Sprague-Dawley rats (timed matings) | Harlan (now Inotiv) | Hsd:Sprague<br>Dawley® SD® |
| **Recombinant DNA** |  |  |
| HA-CD4 | Dr. A.M. Craig | 50 |
| HA-Nrx1β<sup>SS4-</sup> | Dr. T. Sudhof | 51 |
| SEP-GluN1-1a | Dr. R. Malinow | 52 |
| β2m shRNA hairpin | McAllister lab | 45 |
| H2-Kb-mCherry | McAllister lab | <sup>43</sup> |
| NL1-mCherry | This paper | NM_053868.2 |
| pmCherry-N1 | Clontech | 632523 |
| H2-Kb-YFP | This paper | NM_001403363.1 |
| NL1-EGFP | Dr. S. Vicini | <sup>53</sup> |
| **Software and algorithms** |  |  |
| Metamorph | Molecular Devices | V7.5 |
| Prism | GraphPad | V9 |
| MiniAnalysis | Synaptosoft |  |
| **Other** |  |  |
| Neurobasal | ThermoFisher | 21103049 |
| MEM | ThermoFisher | 51200038 |
| Fluoromount | Fisher | 00-4958-02 |
| 18 mm coverslips | Warner | QE-1 |
| Protease/Phosphatase inhibitors | ThermoFisher | A32961 |
| LI-COR Intercept blocking buffer | LI-COR | 927-60001 |

### Resource Availability

#### Lead Contact

Further information and requests for resources and reagents should be directed to and will be fulfilled by the lead contact, Dr. A. Kimberley McAllister.

### Materials availability

This study used two newly generated, unpublished plasmids. These can be made available upon request.

### Data and code availability

- This paper does not report original code
- Any additional information required to re-analyze the data reported in this paper is available from the lead contact upon request.

### Experimental model and subject details

All studies were conducted with approved protocols from the University of California Davis Animal Care and Use Committee, in compliance with NIH guidelines for the care and use of experimental animals. Timed pregnant Sprague-Dawley rats were purchased from Harlan. Mice were generated at UC Davis from C57BL/6, NL1 -/- (knockout; KO), β2m-/- (knockout; KO), and H2-Kb-/- / H2-Db -/- (double KO; DKO) lines. All mouse strains were kept on a pure C57BL/6 background, group housed, and on a 12:12 light/dark cycle. For all experiments, genotypes and treatment were blinded until post-analysis. For all culture experiments, male pups were used. For the β2m-KO biochemistry experiments, three littermate pairs (KO and wild-type; WT) were male and two littermate pairs were female at the indicated age.

### Method Details

#### Reagents

N-methyl-D-aspartic acid (NMDA), glutamate (L-glutamic acid), D(-)-2-amino-5 phosphopentanoic acid (APV), 6-cyano-7-nitroquinoxaline-2,3-dione (CNQX), and α-methyl-4-carboxyphenylglycine (MCPG), all from Sigma-Aldrich (St. Louis, Mo., USA), were dissolved at the indicated concentrations in ACSF (rat) or Neurobasal medium (mouse) from ThermoFisher (Waltham, Ma., USA), with the diluent acting as the control indicated in all figures. All drugs were added 10 min prior to fixation, or following a 10 min control baseline recording via a gravity-fed perfusion system.

#### Neuronal culture and transfections

Neurons from postnatal day (P) 0–2 rat or P1-2 mouse occipital cortex were cultured using established protocols ^43^. Rat neurons were plated at a density of 25K/18 mm coverslip onto either astrocyte monolayers or poly-L-lysine-coated coverslips inverted over astrocyte monolayers as previously described ^43^. Mouse neurons were plated at a density of 45K/18 mm coverslip in astrocyte-conditioned media (AsCM), but otherwise treated the same as rat neurons. AsCM was made by first culturing cortical astrocytes in MEM (Invitrogen, Carlsbad, Ca., USA), media containing 10% fetal calf serum (FCS) (Gemini, Sacramento, Ca., USA) and 1:500 Mycozap (Lonza, Benicia, Ca., USA) and replated after growth to 90% confluency in flasks to standardize astrocyte numbers and remove oligodendrocytes. Upon re-confluence, media was replaced with Neurobasal media supplemented with B27 plus (ThermoFisher, Waltham, Ma., USA), glutamate (ThermoFisher, Waltham, Ma., USA), sodium pyruvate (ThermoFisher, Waltham, Ma., USA), and HEPES (ThermoFisher, Waltham, Ma., USA) for 48 hours, collected, and filter sterilized for storage and use. Rat neurons were transfected with Lipofectamine 2000 (Invitrogen, Carlsbad, Ca., USA) 18-24 hrs before imaging, while mouse neurons were transfected with the same reagent at 3 div for use at 8 div. Data was collected from at least two separate neuronal cultures for all experiments as indicated in the figure legends.

#### Plasmid constructs

HA-CD4 and HA-Nrx1β^SS4−^, lacking an insert in splice site number 4, ^50^ have been described previously ^50,51^. SEP-GluN1-1a cDNA was constructed by fusing the super ecliptic pHluorin (enhanced mutant of pH-sensitive GFP), ^52^ to the N terminus of rat *GluN1-1a*. β2m shRNA hairpin and H2-Kb-mCherry plasmids were previously described ^43,45^. NL1-mCherry (NM_053868.2) was generated by subcloning the rat NL1 sequence into the pmCherry-N1 vector. H2-Kb-YFP was generated by subcloning the mouse H2-Kb sequence into the EYFP-N1 vector. NL1-EGFP was previously described ^53^.

#### Immunocytochemistry

Neurons were fixed with 4% paraformaldehyde and 4% sucrose in PBS at 4°C for 10 min, permeabilized with 0.25% Triton X-100 in PBS for 15 min, blocked with 10% BSA in PBS for 30 min, and incubated with primary and secondary antibodies in 3% BSA. Surface staining was performed on fixed cells without permeabilization. Coverslips were mounted in fluoromount (Fisher, Pittsburgh, PA). Primary antibodies used were as follows: vGlut1 (AB5905, 1:1000; Chemicon, Temecula, CA), GluN2A (07-632, 1:400; Upstate Biotechnology, Lake Placid, NY), GluN2B (AB1557, 1:400; Chemicon), GFP (A11122, 1:1000; Molecular Probes), HA (Clone 3F10, 5ug/ml, Roche), mCherry (AB0040-200, 1:2000; Sicgen), VAMP2 (104-202, 1:1000; Synaptic Systems), and PSD-95 (clone K28/43, 1:750; Antibodies, Inc.). Secondary antibodies used were Alexa Fluor-488, -405, -647, or -568 conjugated anti-rabbit, anti-mouse, and anti-goat (1:200-1:400 Invitrogen).

#### Imaging

Live-imaging was conducted in an imaging perfusion chamber for 18mm coverslips (QE-1; Warner, Hamden, CT) on an Eclipse TE300 Nikon inverted microscope using a 60x oil immersion objective (1.45 NA). Fluorophores were excited at their absorption maxima using a Lambda filter wheel for excitation and emission (Sutter Instrument, Novato, CA) with double and triple band pass filters (Chroma, Battleboro, VT). Bleed-through from fluorophores into other channels was tested by using only one fluorophore and checking all other wavelength and filter combinations for detectable signal. Images were acquired sequentially with a CoolSNAP HQ CCD camera (Roper Scientific, Tucson, AZ) and Simple PCI software (C-Imaging, Compix Inc., Cranberry Township, PA). Imaging was conducted with continual perfusion of ACSF (120 mM NaCl, 3 mM KCl, 2 mM CaCl2, 2 mM MgCl2, 30 mM glucose, 0.2% sorbitol, and 20 mM HEPES, pH 7.3) at 37°C from a gravity-fed perfusion system. Images were typically collected at 10 sec intervals, as this provided the best compromise between high temporal resolution and minimizing neuronal toxicity and photobleaching. For imaging of immunocytochemistry, coverslips were viewed with an Olympus Fluoview 2.1 laser scanning confocal system with a 60x PlanApo oil immersion objective (1.4 NA) on an IX70 inverted microscope. Images for each fluorophore were acquired sequentially with 2.5x digital zoom and 2x Kalman averaging. Rat cortical neuron images used for punctal density or colocalization analysis underwent 1-iteration of deconvolution with the appropriate point spread function for 488, 568 or 647nm light before image analysis (SVI Huygens Essential, Netherlands).

#### Focal Perfusion

Glutamate was perfused locally onto small sections of dendrite through a fire-polished micropipette (tip opening ∼1–2 μm). The micropipette was filled with ACSF containing glutamate (500 µM) and Alexa hydrazide-488 for visualization. Repetitive pressure injection at 2 psi, was applied to the micropipette with an electrically gated valve (Picospritzer, General Valves, Fairfield, NJ) for five, 2 sec pulses at 30 sec intervals for 10 min. For quantification, a region of interest (ROI) was drawn around the size of the focal perfusion area and the mobility of DsRed-GluN1 puncta within the ROI were quantified and compared to non-perfused neighbouring regions of the same dendrite using custom written journals in Metamorph.

#### Electrophysiology

Whole-cell patch-clamp recordings were made from rat cortical neurons at 5-7 div. To isolate mEPSCs, recordings were made at −70 mV in the presence of 0.5 μM TTX. The extracellular solution consisted of 110 mM NaCl, 3 mM KCl, 10 mM HEPES, 10 mM D-glucose, 10 mM glycine, 2 mM CaCl2 and 2 mM MgCl2 at pH 7.3. The osmolarity was adjusted to 305 using sorbitol. The intracellular solution consisted of 130 mM potassium gluconate, 10 mM KCl, 10 mM HEPES, 10 mM EGTA, 1 mM CaCl2, 5 mM ATP-Mg^2+^ and 5 mM GTP-Li^2+^, pH 7.3. Recordings were filtered at 2 kHz using an Axopatch 200B amplifier and digitized at 5 kHZ using Clampex 8 (Axon Instruments). Events were detected with MiniAnalysis software (Synaptosoft), at a threshold four-fold larger than the RMS noise and confirmed by visual inspection. Statistical analysis of average amplitude and frequency was performed using Graphpad Prism software.

#### HEK Cell Co-culture Assay

HEK 293 cells were transfected with lipofectamine-2000 (Invitrogen, Carlsbad, CA) with HA-Neurexin-1β or HA-CD4 plasmids. After 24 hr, transfected cells were trypsinized, plated onto 6 div cortical neurons expressing SEP-GluN1-1a and co-cultured for a further 24 hr. Co-cultured cells were then fixed and immunostained in non-permeabilizing conditions using anti-GFP (A11122, 1:1000; Molecular Probes) and anti-HA (5ug/ml, Roche) antibodies. Confocal Z-stacks were acquired at 0.5 μm steps using a 60x objective on an inverted Olympus Fluoview 2.1 laser scanning confocal system, and processed with MetaMorph software (v7.5, Molecular Devices) using the maximum fluorescent intensity projection. For quantification, the contours of transfected HEK293 cells were chosen as ROIs and the density of dendritic surface GluN1 puncta within the ROI was quantified using custom written journals in MetaMorph.

#### Biochemical experiments

Hippocampi from sex-matched littermate pairs of P15 WT and β2m-KO animals were harvested and dounced in 0.3M Sucrose/Hepes buffer with protease and phosphatase inhibitors (A32961; Thermo). Samples were spun at 600g for 10 min at 4 deg C and the supernatant was collected and denatured in Laemmli buffer with β-mercaptoethanol. Samples were heated at 85 deg C for 15 min and stored at -20 deg C until they were run on SDS-PAGE gel for Western blot analysis. Gels were transferred to nitrocellulose membranes and incubated with primary and secondary antibodies in LI-COR Intercept blocking buffer (LI-COR 927-60001). Secondary antibodies used were donkey anti-rabbit 680 (LI-COR 925-68073) and anti-mouse 800 (LI-COR 925-32212) for imaging on the LI-COR Odyssey system.

#### Data Analysis

Quantification of immunocytochemistry and colocalization during live imaging was performed using the raw images in MetaMorph software ^21,22,27^. Pre- and postsynaptic protein colocalization and density were quantified as described previously using custom journals written in Metamorph software (v7.5, Molecular Devices) ^43^. Intensities were measured within ∼1 µm diameter circles around the centre of manually defined punctae in each channel. Quantification of transport velocities was determined by measuring the time from the start to the end of a unidirectional motion to give mean velocity. Puncta were considered mobile when they performed a unidirectional movement of >2 µm across at least three successive images ^22,27^. Images were processed post-quantification using Adobe Photoshop for presentation purposes.

#### Statistical Analysis

For any experiments with a red dotted line indicated in the figure, conditions were normalized to the control condition within the biological repeat (i.e. individual neuronal cultures) for comparison across experiments. In cases in which multiple genotypes were used, all conditions were normalized to the WT control. For each experiment, the statistical test was selected based on the number of groups being compared, the normality of the distribution, and the equality of the standard deviations. This resulted in Student’s t-tests and ANOVAs as noted in the figure legends. Statistics were run using Graphpad Prism (V9) software. All data are expressed as mean ± SEM.

## Acknowledgements

Conflict of Interest: The authors declare no competing financial interests. This work was supported by NIH R01-EY13584 (A.K.M.) and R01-NS060125 (A.K.M.) and by philanthropic support from H. Britton Sanderford, Jr. We thank Ann-Marie Craig for providing HA-CD4, Thomas Sudhof for providing HA-Nrx1β^SS4−^, and Dr. R. Malinow for providing SEP-GluN2A. The authors would also like to thank Drs. Samantha Spangler and Leigh Needleman for generating rat neuronal cultures and Faten El-Sabeawy for contributing valuable technical support.

## Results

### Acute neuronal activity alters glutamatergic synapse density within minutes

At very early stages of CNS development, even prior to synapse formation, dendrites are speckled with highly mobile ionotropic glutamate receptor transport packets that can detect, and be activated by, ambient glutamate as they are rapidly cycle along the dendritic shaft ^21,27^. Given that their recruitment to nascent synapses occurs within approximately 8 min following initial axonal contact ^21^ and activity regulation as short as 5 min significantly alters the phosphorylated proteome in synapses ^54^, it is conceivable that neurotransmitter release during cortical development could not only regulate cortical connectivity and the number of synapses formed but also could do so in a timeframe of minutes rather than hours or days as previously reported ^55,56^. To test this hypothesis, neurotransmitter inhibitors and agonists were applied to 6-7 div rat cortical cultures 10 min prior to fixing and immunostaining with antibodies against the presynaptic vesicle protein vGlut1 and the postsynaptic NMDAR subunits GluN2A and GluN2B to label synapses. The inhibitors used were TTX (500 nM) to block action potentials, CNQX (10 µM) to block α-amino-3-hydroxy-5-methyl-4-isoxazolepropionic acid receptor (AMPARs), and APV (50 µM) to block NMDARs, as well as a receptor antagonist cocktail (ACM) comprised of APV, CNQX, and MCPG (500 µM) to also block metabotropic glutamate receptors. We found that, after just 10 min of APV treatment, there was a significant 25% increase in the density of synapses determined by quantification of overlap between vGlut1 and GluN2A/GluN2B (**Figure 1A, B**), whereas no changes were observed following TTX, CNQX or ACM treatment (**Figure 1B**). Conversely, stimulation of NMDARs for just 10 min with either NMDA (50 µM), or NMDA/Glycine (N/Gly, 1 mM/10 µM) or glutamate (50 µM) all elicited a significant decrease in glutamatergic synapse density by 23, 30 and 35% respectively (**Figure 1B**). Spontaneous activity is developmentally regulated in cortical sensory areas ^57^, so we next determined if activity regulates synapse density in a similar way at later ages. Neither stimulating with glutamate or activators of NMDARs, or blocking with ACM or APV elicited changes in synapse density in older cultures (14 div, **Figure 1C, D**). Thus, activation of NMDARs rapidly and negatively alters glutamatergic synapse density selectively during the period of initial establishment of cortical synapses.

**Figure 1.**
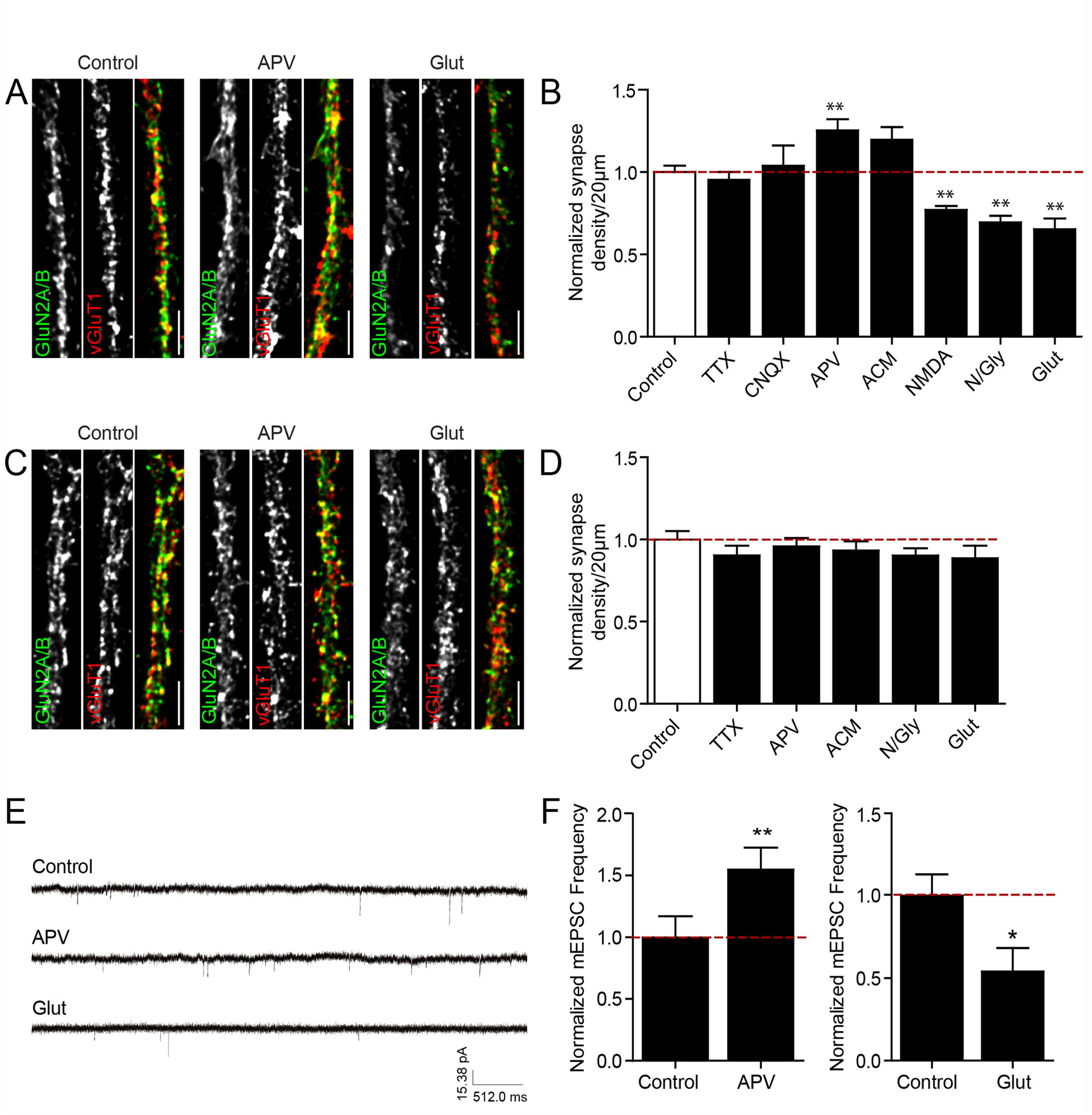
Activity rapidly regulates cortical synapse density in young neurons (A) Representative confocal images of dendrites show the density of synapses, determined by colocalization between postsynaptic GluN2A and GluN2B puncta (*green*) and presynaptic vGlut1 puncta (*red*), from cortical neurons at 6-7 div incubated with ACSF (control; left panels), APV (middle panels), or glutamate (Glut: right panels) for 10 min before fixation. (B) Glutamatergic synapse density was significantly increased following blockade of NMDARs with APV for 10 min and decreased following activation of NMDARs with NMDA, NMDA plus glycine (N/Gly) or Glut. In contrast, a cocktail of APV, CNQX and MCPG (ACM), or TTX or CNQX treatment alone did not change synapse density compared to control. Control n = 20, TTX = 13, CNQX = 8, APV = 24, ACM = 8, NMDA = 22, N/Gly = 14, Glut = 9 cells. ** p < 0.01, ANOVA. (C) Representative confocal images of dendrites from 14 div cortical neurons following 10 min treatment with ACSF (control), APV, or glutamate. (D) Glutamatergic synapse density was unaltered following acute blockade or activation of NMDARs for 10 min in older cultures. Control n = 9, TTX = 8, APV = 9, ACM = 12, N/Gly = 10, Glut = 8 cells. ANOVA. (E) Representative traces of mEPSCs from 7 div cortical neurons incubated for 10 min with ACSF (control), APV, or glutamate. (F) Acute blockade of NMDARs with APV significantly increased mEPSC frequency, whereas acute activation of NMDARs with glutamate significantly decreased it. Control n = 13,14, APV = 16, glutamate = 13 cells. * p < 0.05, ** p < 0.01, t-test. Data is plotted as mean +/- SEM, normalized to control cells from the same culture/experiment (*red dotted line*). Scale bar, 5µm.

Since excess glutamate can be excitotoxic to neurons, we next examined whether acute application of glutamate induced cell death under our experimental conditions (**Supp. Fig. 1**). 6 div cortical neurons, expressing GluN1-DsRed, were treated with glutamate (50 µM) for 10 min and then loaded with the cell death fluorescent marker Sytox Green (0.5 µM). This DNA-binding dye stains the nuclei of cells with a compromised plasma membrane, and therefore represents an ideal indicator of cell death ^58^. Following a 10 min incubation of cells with glutamate, no detectable Sytox Green fluorescence was observed. Neuronal health was confirmed by examining the neurons under brightfield illumination. Neurons were round and translucent in appearance, characteristic of healthy viable cells (**Supp. Fig. 1**).

Next, to determine if manipulation of NMDAR activation alters the number of functional synapses, in addition to its effects on synapse density, whole-cell patch clamp recording of spontaneous miniature excitatory postsynaptic currents (mEPSCs) was performed. Following blockade of NMDARs for 10 min with APV, mEPSC frequency increased by 55% on average, compared to control neurons incubated with ACSF (**Figure 1E, F**). Conversely, and with a similar bidirectional result to synapse density measured with ICC, glutamate stimulation induced a significant reduction in mEPSC frequency by 46% (**Figure 1E, F**). There was no change in mEPSC amplitude under any condition (*not shown*). Taken together, these results provide evidence that during the initial stages of synapse formation, glutamate release can rapidly, within minutes, alter the density of synapses on cortical neurons.

### Glutamate rapidly alters NMDAR transport and surface expression

Because synapse formation requires transport of NMDARs in NRTPs to sites of contact ^21^, it is possible that the rapid glutamate-induced decrease in synapse density could be due to alterations in NRTP transport (**Figure 2A-H**). To test this hypothesis, young cortical neurons at 4 div were transfected with GluN1-DsRed and time-lapse imaged 24 hr later, 10 min before and 10 min during ACSF, ACM, or glutamate treatment (**Figure 2A-C**). The mobile fraction of NRTPs, calculated as the percent of total GluN1DsRd puncta that moved >2 µm during the imaging period, was substantially altered following 10 min incubation with either glutamate receptor antagonists or agonists. Acute incubation with APV for 10 min increased NRTP mobility by roughly 29% compared to that of control ACSF treated neurons (**Figure 2G**), while ACM elicited a significant increase in the number of mobile NRTPs by approximately 64% (**Figure 2B, G**). Conversely, acute glutamate exposure significantly decreased the proportion of mobile NRTPs to less than 50% of control (**Figure 2C, G**), as did treatment with NMDA and N/Gly, which reduced mobility by 45 and 61% respectively (**Figure 2G**). In fact, the effects of glutamate were so striking that cessation of NRTPs occurred almost instantaneously following the addition of glutamate (**Figure 2C**), suggesting that mobile NRTPs can sense and fully respond to ambient glutamate almost immediately ^27^.

**Figure 2.**
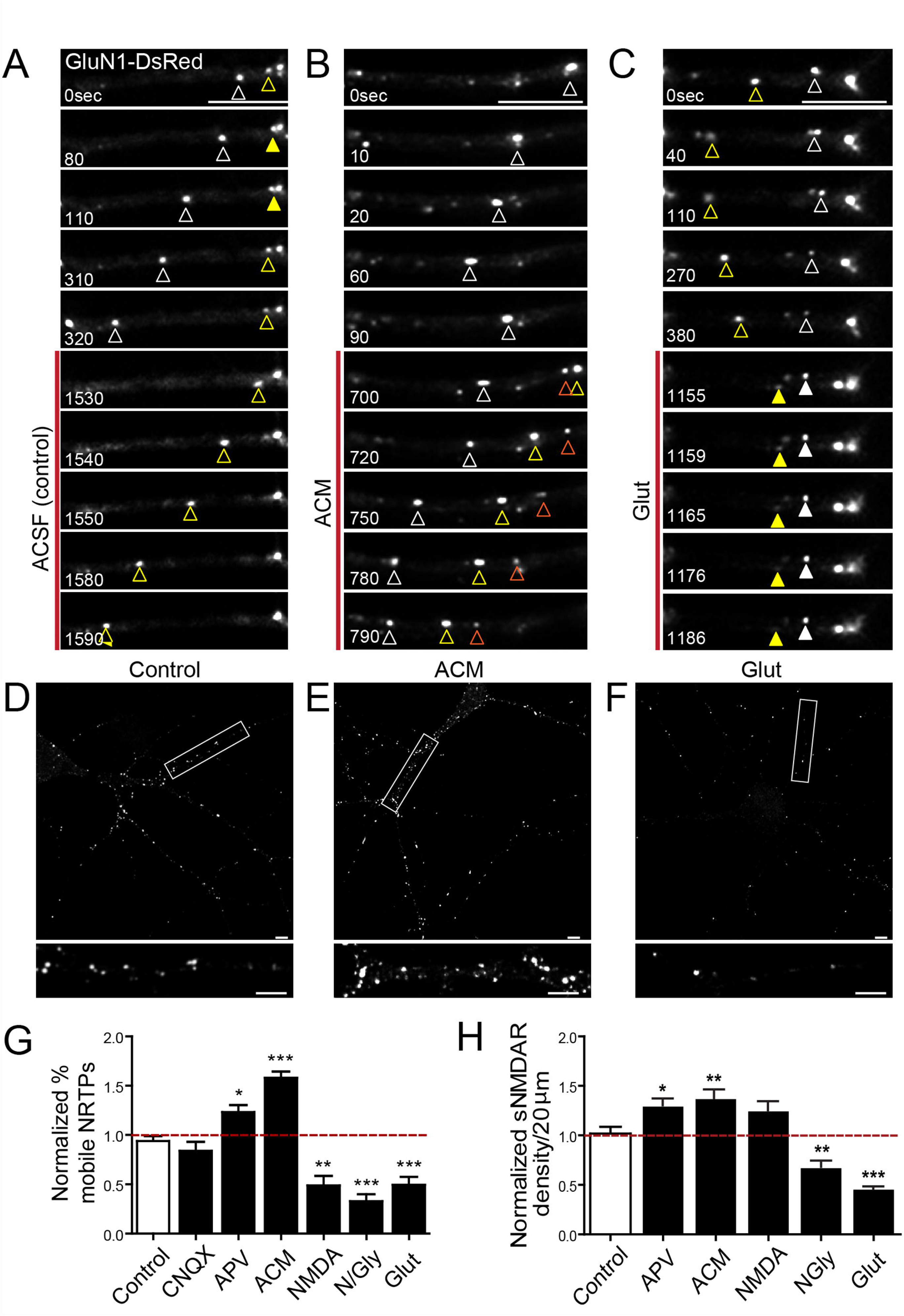
NMDAR activation rapidly regulates NRTP transport (A-C) Time-lapse images of dendrites from GluN1-DsRed-expressing 5 div cortical neurons before and during acute (10 min) treatment (indicated by the red line) with (A) ACSF (control), (B) a cocktail of glutamate receptor inhibitors (ACM) or (C) glutamate. Arrowheads indicate mobile (*open arrowheads*) and stable (*closed arrowheads*) NRTPs moving throughout the dendrite. Time is in sec. (D-F) Representative confocal images of 5 div cortical neurons expressing SEP-GluN1-1a, fixed and stained with antibodies against GFP in non-permeabilizing conditions, following 10 min treatment with (D) ACSF (control), (E) ACM or (F) glutamate. Boxed dendritic regions are magnified below each respective image. (G) Blockade of NMDARs following 10 min incubation with APV or ACM significantly increased NRTP mobility, whereas acute activation of NMDARs with NMDA, N/Gly and glutamate significantly decreased it. All values are shown normalized to NRTP mobility in the same dendritic segments prior to drug treatment (*red dotted line*). Control n = 8, CNQX = 10, APV = 9, ACM = 8, NMDA = 5, N/Gly = 7, Glut = 9 cells. (H) Acute activation of NMDARs with APV and ACM significantly increased surface NMDAR (sNMDAR) expression, while N/Gly and glutamate significantly decreased it. Values are shown normalized to control (*red dotted line*). Control n = 17, APV = 16, ACM = 14, NMDA = 5, N/Gly = 15, Glut = 11 cells. Data is plotted as mean +/- SEM, normalized as indicated for each panel. * p<0.05, ** p<0.01, *p<0.001, ANOVA. Scale bar, 5µm.

Another measure of receptor trafficking is their surface expression. We therefore tested whether glutamate treatment and NMDAR activation altered the number of NMDARs expressed on the surface of the plasma membrane. To reliably detect surface NMDARs, an N-terminal tagged GluN1 SEP construct (SEP-GluN1-1a) was used so that we could detect the SEP tag on the surface of the cell using antibodies against GFP under non-permeabilized staining conditions ^59^. 8 div cortical neurons that had been transfected with SEP-GluN1-1a at 5 div were acutely exposed to activators or inhibitors of NMDARs for 10 min and then immediately fixed and stained, in non-permeabilizing conditions, with antibodies to GFP (**Figure 2 D-F**). Consistent with the bi-directional changes observed for synapse density and NRTP mobility, blockade of NMDARs with APV or with the antagonist cocktail significantly increased the proportion of total NMDARs on the dendritic surface by 28% and 37%, respectively, while glutamate and N/Gly treatment significantly decreased it by 55 and 34% respectively (**Figure 2H**). Taken together, these results strongly suggest that activity may rapidly regulate synapse density in young neurons via rapid changes in NMDAR mobility and surface expression.

### Glutamate-induced alterations in NRTP transport require Ca^2+^ influx through NMDARs

NMDARs are cation channels, permeable to both sodium (Na^+^) and Ca^2+^ ions. NMDAR-mediated influx of Ca^2+^ initiates a second messenger cascade responsible for regulating a diverse array of neuronal functions during development including neurite outgrowth, and long-term potentiation (LTP) and long-term depression (LTD) ^60-63^. We therefore determined whether alterations in NRTP mobility following glutamate treatment were dependent on Ca^2+^ influx following NMDAR activation. First, we imaged NMDAR-mediated Ca^2+^ influx elicited by glutamate using the high-affinity cell-permeant Ca^2+^ indicator Fluo-4 AM in cortical neurons expressing GluN1-DsRed to simultaneously monitor NRTP transport and changes in intracellular Ca^2+^. 4 div cortical neurons were transfected with GluN1-DsRed, then loaded with Fluo-4 24 hr later. Time-lapse imaging was then performed 10 min prior to and during acute application of the ACM inhibitor cocktail (**Figure 3A**) or glutamate (**Figure 3B**). Upon addition of ACM to prevent NMDAR activation and Ca^2+^ influx, NRTP mobility increased, as expected, while no changes in Fluo-4 fluorescence were observed (**Figure 3A, C**). In contrast, glutamate elicited a sustained intracellular Ca^2+^ rise that remained elevated throughout the 10 min imaging period. Simultaneous monitoring of NRTP mobility revealed that this Ca^2+^ elevation occurred immediately prior to cessation of mobility (**Figure 3B, D**). To further confirm that Ca^2+^ influx was responsible for inhibition of NRTP mobility following glutamate stimulation, neurons were pre-incubated with BAPTA-AM, a Ca^2+^ chelator. In the presence of BAPTA, NRTP mobility remained unchanged following acute glutamate exposure (**Figure 3E**) compared to the 67% reduction in mobility observed with glutamate. In addition to Ca^2+^ influx, the ability of glutamate to alter NRTP transport also depended on its ability to bind to the receptor, since incubation with glutamate in the presence of APV to block NMDARs, fully prevented glutamate-induced inhibition of NRTP transport. CNQX had no effect, suggesting that influx through AMPARs was not required (**Figure 3E**). Thus, glutamate requires Ca^2+^ influx through NMDARs to rapidly decrease NRTP transport.

**Figure 3.**
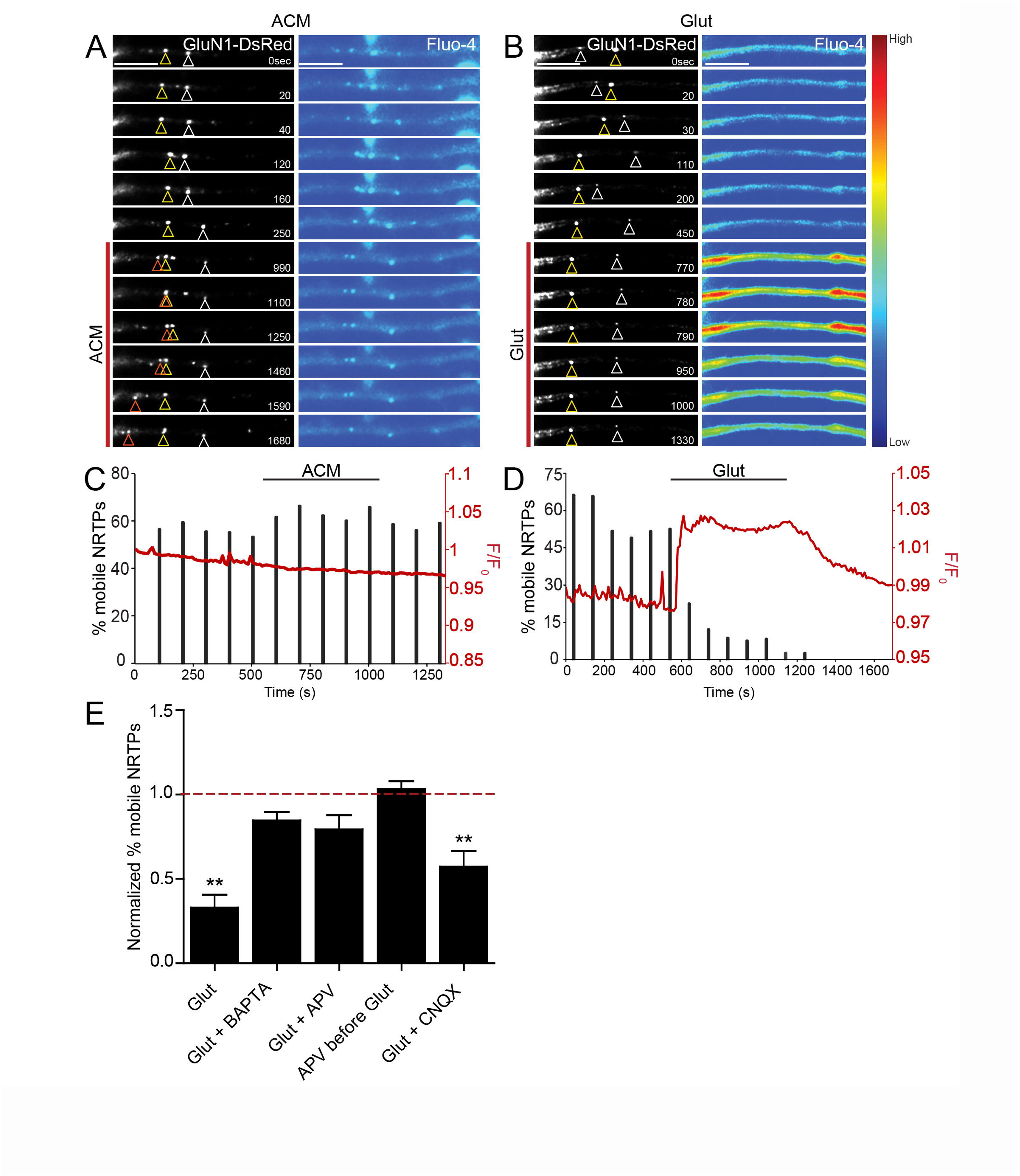
The effects of glutamate on NRTP mobility are NMDAR-dependent and require Ca^2+^ influx (A and B) Representative time-lapse images of 5 div cortical neurons expressing GluN1-DsRed (*left column*) and loaded with Fluo-4 AM (*right column*), to simultaneously monitor NRTP mobility and intracellular Ca^2+^ in the same cell, before and during 10 min ACM (A) or glutamate (B) treatment indicated by the red line. The magnitude of the changes in Fluo-4 signal are shown in pseudocolor. Scale bar, 5µm. (C and D) The increase in NRTP mobility (*black bars*) following acute exposure to ACM (C) occurred in the absence of an increase in Ca^2+^ (F/F0, *red trace*) whereas the inhibition of NRTP mobility following acute treatment with glutamate (D) occurred immediately following influx of Ca^2+^ through NMDARs. ACM n = 6, Glut = 5 cells. Note the timescale is extended post-glutamate in D to monitor whether the glutamate effect washed out. (E) BAPTA-AM, and APV treatment prevented the glutamate-induced reduction in NRTP mobility, whereas CNQX was without effect. In the APV before Glut condition, APV was added 2 minutes prior to glutamate addition. Glut n = 4, Glut + BAPTA =5, Glut + APV = 5, APV before Glut = 7, Glut + CNQX = 6 dendrites. Data is plotted as mean +/- SEM, normalized to GluN1 mobility in the same dendritic segments prior to drug treatment (*red dotted line*). **p<0.01, t-test. Unless indicated, drug concentrations are as in Figure 1.

### Glutamate acts locally to alter NRTP transport

We next tested whether local changes in glutamate, as may occur when axons approach dendrites during synapse formation, also alter NRTP transport. Using focal perfusion, pipettes, pulled to a tip diameter of 1-2 µm, were positioned about 10 µm from a dendritic segment expressing GluN1-DsRed. A suction pipette was positioned upstream of the application pipette, enabling a small, confined stream of glutamate or ACSF to be locally applied to a limited section of dendrite, while avoiding other dendrites or the cell soma (**Supp. Fig. 2A-C**). ACSF (**Suppl. Fig. 2A**), glutamate (500 µM) (**Supp. Fig. 2B**), or glutamate in the presence of APV (**Supp. Fig. 2C**) was applied focally by pressure application (2 psi) for five 2 sec pulses at 30 sec intervals for 10 min. For each experiment, neighbouring, non-focally perfused regions of the same dendrite were used for comparison; data was normalized to pre-treatment NRTP mobility (**Supp. Fig. 2D**).

Focal perfusion of ACSF did not itself alter NRTP trafficking since NMDARs moved freely, exhibiting normal mobility and bi-directional transport behaviour throughout the focally-perfused segment of dendrite (**Supp. Fig. 2A)**. However, perfusion of glutamate onto limited sections of dendrites induced local inhibition of NRTP mobility by 35% specifically within the perfused region, compared to neighbouring, non-focally perfused regions (**Supp. Fig. 2B, D**). APV (500 µM), when perfused together with glutamate, prevented the inhibitory effect of glutamate on NRTP mobility, presumably by blocking NMDAR-mediated Ca^2+^ influx; NRTPs continued to travel bidirectionally throughout the dendrite in this condition (**Supp. Fig. 2C, D**). Finally, NMDARs in the neighbouring non-perfused regions were not attracted to glutamate since there were no changes in directionality towards or away from the perfused region (**Supp. Fig. 2E**). Thus, activity can act locally to rapidly alter NRTP dynamics prior to synaptogenesis.

### Glutamate inhibits trafficking of NL1 and its cotransport with NRTPs

The cotransport of NRTPs with NL1 is critical for the rapid recruitment of NMDARs to nascent synapses as they form during development ^22,64^. Given that glutamate inhibits NRTP mobility, we hypothesized that glutamate would also decrease the cotransport of NRTPs with NL1 and potentially even their colocalization. To test this idea, 4-5 div cortical neurons were co-transfected with GluN1-DsRed and eGFP-NL1 and time-lapse imaged 24 hr later to visualize the dynamics of NRTPs and NL1 puncta simultaneously. NRTP and NL1 puncta exhibited dynamic behaviour, rapidly moving bi-directionally throughout dendritic compartments (**Figure 4A-B**), as previously reported ^21,22,27^. In addition to rapid inhibition of NRTP mobility, acute application of glutamate (50 µM) dramatically and rapidly decreased NL1 mobility (**Figure 4C**) and caused an 87% reduction in NRTPs co-transported with NL1 (**Figure 4D**) without significantly altering overall NL1/GluN1 colocalization (**Figure 4E**). Similarly, 10 min treatment with N/Gly significantly reduced the co-transport of NRTPs with NL1 by 74% (**Figure 4D**). Thus, the inhibitory effect of glutamate is not restricted to NRTPs, but also rapidly alters the transport of other synaptic proteins involved in synaptogenesis. The loss in mobility between colocalized NL1 and NRTPs could explain the rapid loss in synapse density observed following acute glutamate treatment. In contrast, bath application of the antagonist cocktail, ACM, failed to have any effect on NL1 mobility or its colocalization or co-transport with NRTPs, suggesting that the population of NRTPs that exhibit a rapid increased mobility following ACM treatment (**Figure 2G**) are likely not associated with NL1 (**Figure 4C-E**). These results demonstrate that rapid alterations in glutamate release can specifically disrupt the cotransport of NRTPs and NL1, thereby preventing their recruitment or stabilization at nascent synapses.

**Figure 4.**
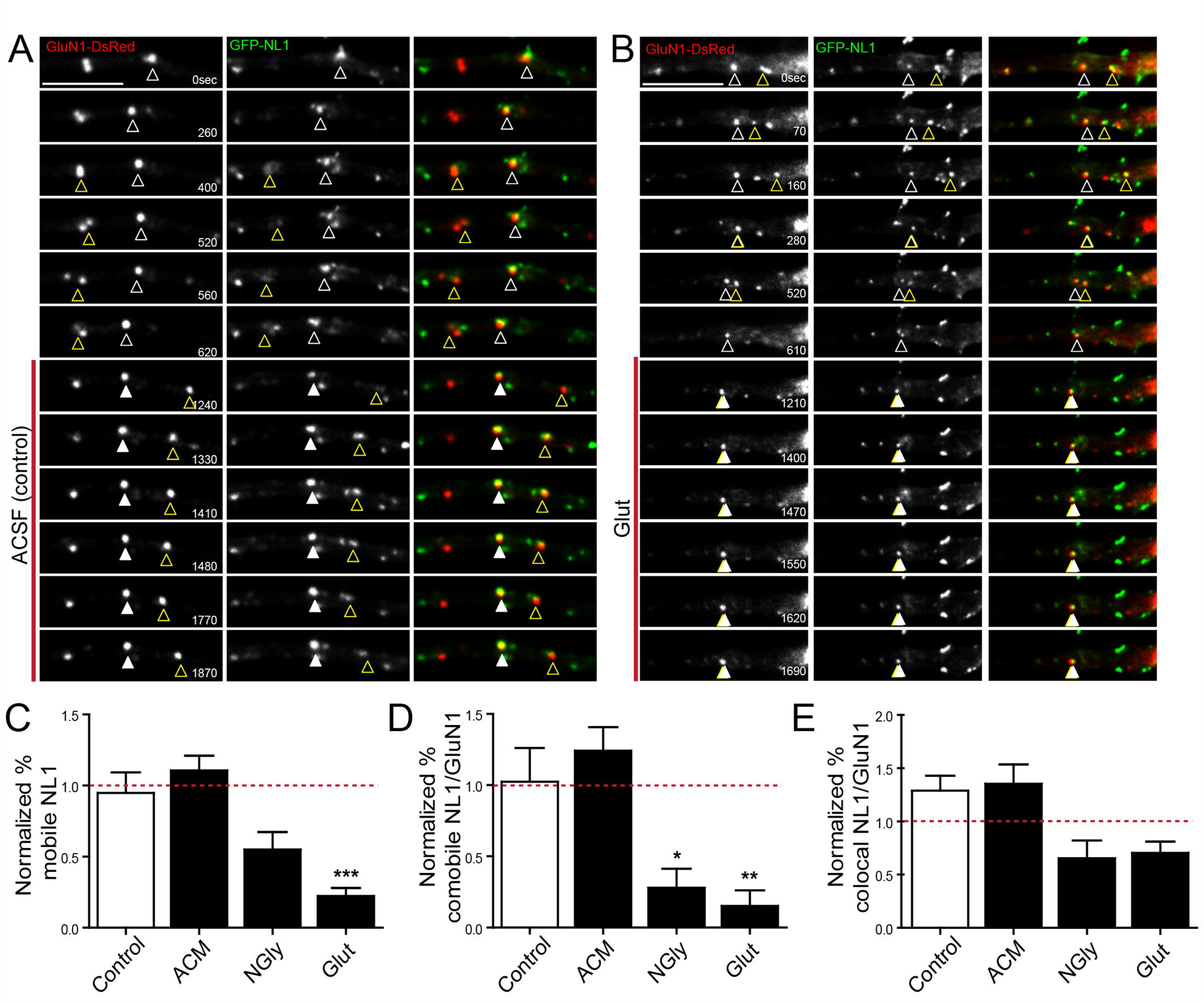
Glutamate inhibits the transport of NL1 and its cotransport with NRTPs. (A and B) Time-lapse images of 5 div cortical neurons simultaneously expressing GluN1-DsRed (*left column*) and GFP-NL1 (*central column*) to visualize co-localization (*overlay, right column*) and co-transport, before and during (*red line*) ACSF (A) or glutamate (B) treatment. Open and closed arrowheads indicate mobile and stationary puncta respectively. Scale bar, 5µm. (C) Acute glutamate treatment, but not ACM or NMDA/glycine significantly decreased NL1 mobility. (D) Acute glutamate and NMDA/Glycine treatment, but not ACM, significantly decreased NL1/GluN1 co-mobility. (E) NL1 colocalization with GluN1 was not significantly altered by any of the treatments. The same samples were analysed for graphs in (C-E): ACSF n = 6, ACM = 15, NGly = 9, Glut = 9. Data is plotted as mean +/- SEM. Data was normalized to measures in same dendritic segment before treatment (*red dotted line*). *p<0.05, **p<0.01, ***p<0.001, ANOVA.

### Decreased NL1 is necessary for glutamate-induced synapse loss

To determine if acute glutamate-induced synapse loss requires changes in NL1 levels, we switched our assay to mouse neurons so we could use a genetic deletion of NL1 ^47,65^. In our hands, mouse cultures tend to have lower synaptic density compared to the rat cultures used in earlier figures. First, we confirmed that 8 div occipital WT mouse neurons show the glutamate-induced reduction in excitatory synapses (**Supplemental Figure 3A-B**). Next, we tested whether acute glutamate treatment requires a decrease in NL1 to cause synapse loss, since NL1 is well-known to stabilize synapses at older ages ^16^. If so, then preventing the glutamate-induced decrease in NL1 should prevent synapse loss. Cortical neurons were transfected with a construct to overexpress NL1 (NL1-mCherry) at 3 div, treated with or without glutamate for 10min at 8 div, and synapses were quantified as above. Consistent with our hypothesis, preventing the decrease in NL1 by overexpression of NL1-mCherry rescued the glutamate-induced synapse loss (**Figure 5A-B**), suggesting that excess NL1 stabilizes a higher proportion of excitatory synapses to offset the effects of glutamate. Thus, decreases in NL1 levels are necessary for the effects of glutamate in causing synapse loss.

**Figure 5.**
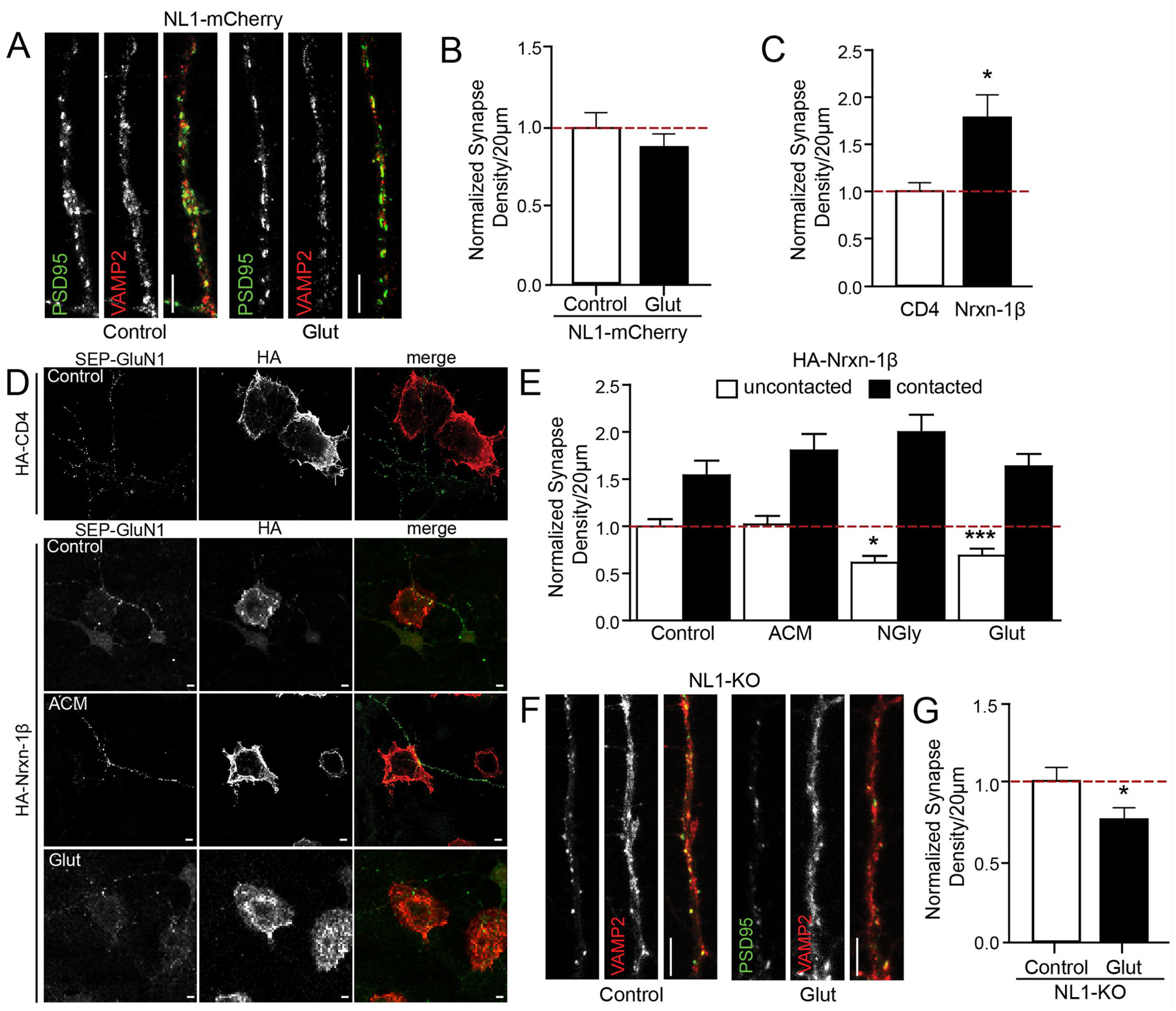
NL1/Nrxn binding prevents glutamate-induced internalization of sNMDARs and synapse loss (A) Representative confocal images of occipital WT cortical mouse neurons transfected with NL1-mCherry that were incubated with glutamate or control for 10 min at 8 div, then fixed and stained with antibodies against VAMP2 and PSD-95. Synapses were defined as colocalized postsynaptic PSD95 (*green*) and presynaptic VAMP2 puncta (*red*). (B) Glutamatergic synapse density was not altered following 10 min glutamate treatment of occipital WT mouse cultures. Data is plotted as mean +/- SEM, normalized to the values from control cells from the same culture/experiment (*red dotted line*). Control n = 26, glut = 23 cells, t-test. (C) Density of sGluN1 puncta was calculated for dendritic segments contacting HA-positive HEK cells expressing HA-CD4 (*open bar*) or HA-Nrxn-1β (*black bar*) normalized to the HA-CD4 condition (*red dotted line*). HA-Nrxn-1β contacting dendrites showed a significantly increased density of sGluN1 puncta. HA-CD4 n = 12, HA-Nrxn-1β n = 6 dendrites, * p < 0.05, t-test. (D) Representative maximum projection images of HEK cells expressing HA-CD4 (control) or HA-Nrxn-1β co-cultured with cortical neurons at 6 div expressing SEP-GluN1-1a, fixed and immunostained for GFP (*green*) and HA (*red*) following 10 min treatment with ACM or glutamate. Scale bar, 5µm. (E) Glutamate and NMDA/glycine, but not ACSF or ACM, treatment caused a loss of surface GluN1 in sections of dendrites that were not in contact with HA-Nrxn-1β expressing HEK cells (*open bars*). This glutamate-induced loss is prevented at contacts with HEK cells expressing HA-Nrxn-1β (*black bars*). Density is normalized to sGluN1 density in non-contacting dendrites from vehicle treated cells (control) (*red dotted line*). Control n = 16, ACM n = 23, NGly = 14, Glut = 23 cells. * p < 0.05, *** p < 0.001, ANOVA. (F) Representative confocal images of occipital NL1-KO cortical mouse neurons treated with glutamate and stained for PSD95 (*green*) and VAMP2 puncta (*red*). Scale bar, 5µm. (G) Glutamatergic synapse density in NL1-KO cultures was significantly decreased following 10 min of glutamate treatment. Data is plotted as mean +/- SEM, normalized to the control cells treated with vehicle from the same culture/experiment (*red dotted line*). NL1-KO control n = 42, NL1-KO glut = 27 cells, *p<0.05, t-Test.

### Nxrn/NL1 adhesion counteracts the inhibitory effects of glutamate

So far, our results suggest that NL1 opposes the effects of glutamate in causing synapse loss and NMDAR internalization. This model is similar to the role for ACh at the NMJ where ACh destabilizes postsynaptic sites through the dispersal of ACh receptors (AChRs) ^38^. This effect is counteracted by an anti-dispersal signal in the form of agrin, which prevents neurotransmitter-induced AChR loss^38^. Since NL1 mediates the trafficking and recruitment of NMDARs to nascent synapses ^22,53^, and since it requires binding to its presynaptic ligand Nrxn1 in order to initiate synapse formation ^66-69^, we reasoned that NL1 and its interaction with Nrxn1 may act as a similar anti-dispersal signal to protect NMDARs against the effects of glutamate.

To test this hypothesis, we utilized a hemi-synapse assay ^70^. Cortical neurons at 6 div were cocultured with HEK293 cells expressing HA-Nrxn-1β in the presence or absence of glutamate or N/Gly to activate, or ACM to block, NMDARs, respectively. In this classic hemi-synapse assay, postsynaptic specializations are induced to cluster along dendrites at sites of contact with co-cultured HEK cells expressing the synaptogenic molecule Nrxn1 ^70,71^. We used the splice variant of Nrxn1β lacking an insert at splice site 4 that potently binds with postsynaptic NL1 ^50^. HA-Nrxn-1β-induced clustering of postsynaptic GluN1 was assessed using antibodies to GFP to detect surface-expressed SEP-GluN1-1a, and to HA to detect CD4- or Nrxn1-expressing HEK cells. As expected ^70,71^, GluN1 was induced to cluster selectively at sites of contact with Nrxn1-expressing HEK cells, compared to HEK cells expressing HA-CD4 control (**Figure 5C-D**). Also, as expected from data presented above, acute incubation with glutamate caused a 31% loss in surface GluN1 puncta in dendritic segments that were not in contact with HEK cells. Importantly, this glutamate-induced loss of GluN1 was prevented at sites of contact with Nrxn1-expressing HEK cells. Similarly, the 38% loss of surface GluN1 induced by N/Gly was also prevented in Nrxn1-contacting dendrites compared to the non-contacting dendrites (**Figure 5E**). Thus, binding of NL1 to Nrxn1 stabilizes the complex and protects NMDARs from dispersal by glutamate, similar to the role for agrin as an anti-dispersal signal that protects AChRs from ACh-induced dispersal at the NMJ ^38^.

Finally, if our hypothesis is true, then glutamate should still cause synapse loss in neurons lacking NL1 because of the lack of this anti-dispersal signal. Indeed, 10 min of glutamate treatment induced a significant 25% reduction in excitatory synapses in 8 div NL1-KO neurons compared to control NL1-KO neurons (**Figure 5F-G**). Thus, Nrxn-NL1-induced NMDAR puncta are resistant to glutamate-induced loss, suggesting that an interaction across the synaptic cleft protects and stabilizes developing synapses from the effect of glutamate release. Together, these results indicate that NL1 is a critical component of the anti-dispersal signal that opposes glutamate-induced synapse loss in cortical neurons during the early stages of synapse formation.

### MHCI is necessary for glutamate-induced synapse loss through negatively regulating NL1

To further dissect the molecular mechanisms that mediate the effects of glutamate on synapse density, we hypothesized that other molecules present at the synapse that cause synapse loss in an activity-dependent manner may play a role. MHCI molecules are optimal candidates since they are activity-regulated, expressed in neurons at synapses ^44,72,73^, and negatively regulate synapse density selectively during the period of the initial establishment of connections when glutamate rapidly alters synapse number ^43,45^. Importantly, little is known about how MHCI causes synapse loss^17,74^.

If glutamate requires MHCI to cause synapse loss, then we would expect that acute glutamate treatment would increase MHCI levels in neurons and especially surface MHCI (sMHCI) which is required for synapse loss ^43^. WT occipital 8 div cultures were treated with 50μM glutamate and sMHCI levels were quantified. While total MHCI levels are clearly regulated by activity over longer time-frames of days ^43^, acute effects of altering neural activity on MHCI levels are unknown. Although 10min of glutamate did not significantly alter sMHCI density (**Supplemental Figure 3C-D**), it did significantly increase the intensity of sMHCI puncta (**Figure 6A-B**). Thus, MHCI levels are increased by acute glutamate treatment.

**Figure 6.**
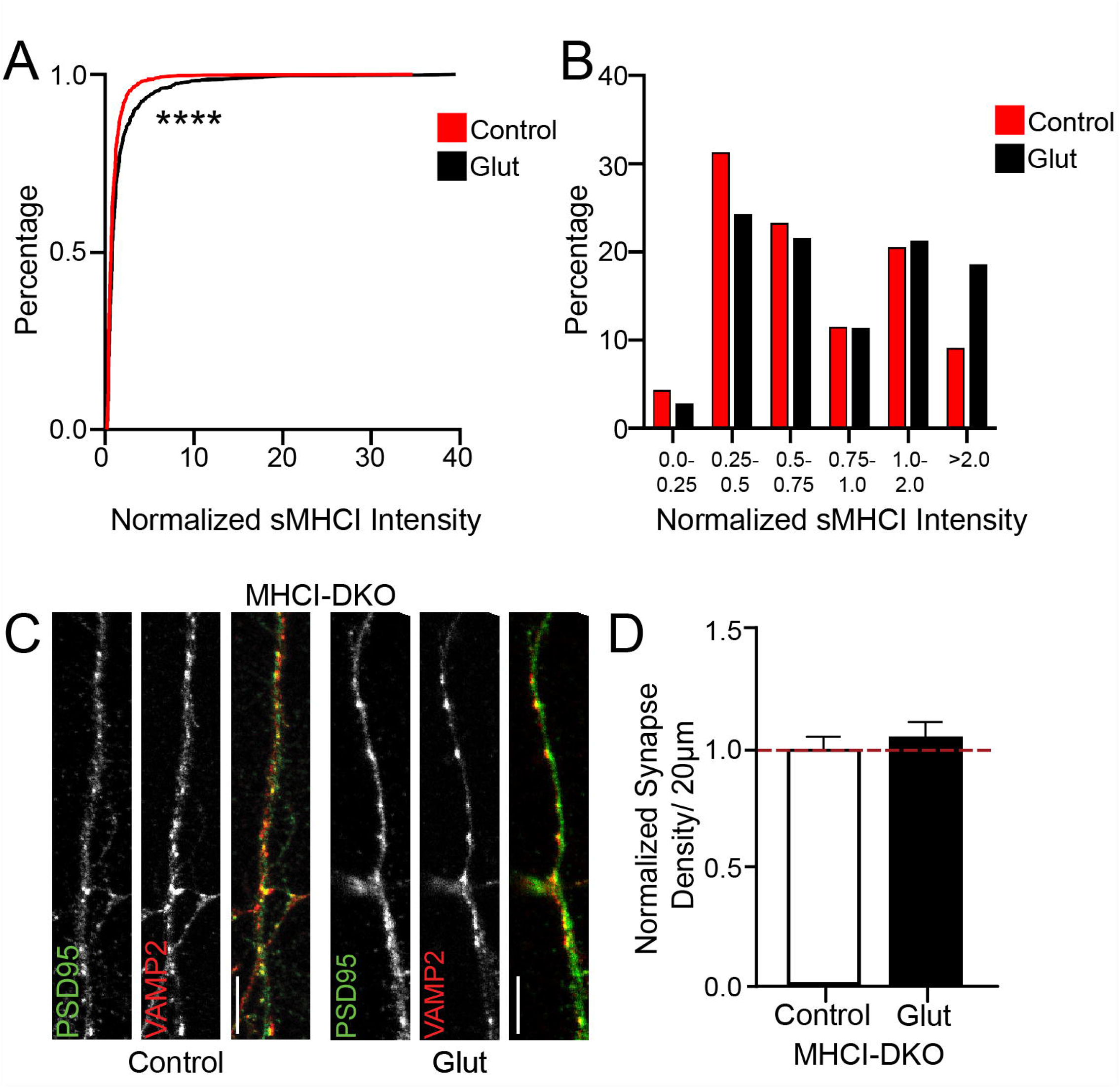
An increase in surface MHCI is required for glutamate-induced synapse loss (A) sMHCI intensity shifts to more intense puncta in response to glutamate treatment in WT DIV8 occipital neurons. Control n = 21 cells, 922 puncta, glutamate n = 27 cells, 1158 puncta. Kolmogorov-Smirnov test, ****p<0.0001. Data is plotted as a cumulative probability plot for control (*red*) and glutamate (*black*), with intensity normalized to control cells from the same culture/experiment. Specifically, intensity for sMHCI was averaged for each neuron in each of three independent cultures. From that average, intensity was normalized within culture, thus reflecting a control average of “1”. (B) MHCI punctal intensity was binned and graphed as a percentage of the whole for both control and glutamate. There is a shift to higher intensity puncta in the glutamate, most clearly illustrated in the decrease in the proportion of puncta with intensity between 0.25-0.5 and an increase in intensity in puncta with intensity greater than 2. (C) Representative confocal images of MHCI-DKO occipital cortical mouse neurons at 8 div that were incubated with glutamate for 10 min, then fixed and stained with antibodies to postsynaptic PSD95 (*green*) and presynaptic VAMP2 puncta (*red*); glutamatergic synapses are defined as colocalized puncta. Scale bar, 5µm. (D) Glutamatergic synapses were not significantly altered by glutamate treatment in the absence of H2-Kb and H2-Db. MHCI-DKO control n = 29, MHCI-DKO glut = 28 cells, t-test.

Our lab has previously demonstrated that elevations in MHCI cause a loss of NMDARs and glutamatergic synapses selectively during the same early developmental period that glutamate rapidly alters synapses ^43,45^. To test whether increases in MHCI are necessary for glutamate-induced synapse loss, 8 div neurons from mice that lack the two major classical forms of MHCI, H2-Kb and H2-Db (MHCI-DKO) were treated with glutamate or control media for 10min before fixing and staining for PSD-95 and VAMP2 (**Figure 6C**). Glutamate failed to reduce synapse density in neurons that lacked H2-Kb and H2-Db (**Figure 6D**), showing the necessity of MHCI for glutamate-induced synapse loss in early cortical cultures.

Because both MHCI and NL1 are required for glutamate-induced synapse loss, we next asked if MHCI acts upstream of NL1 to mediate glutamate-induced changes in synapse density. First, we determined if altering MHCI levels negatively regulate NL1 using cultured cortical neurons and a lentiviral strategy to overexpress MHCI H2-Kb-YFP from 3-8 div, with GFP or NL1-shRNA as a negative and positive control, respectively. When measured via Western blot and normalized to actin as a loading control, H2-Kb-YFP significantly decreased NL1 protein levels by 34% compared to GFP expression alone (**Figure 7A-B**). Using the β2m-KO mouse in which most forms of classical MHCI are not expressed on the surface of cells ^44^, we harvested and lysed sex-matched littermate hippocampi for Western blot analysis (**Figure 7C**). Removal of surface MHCI significantly increased NL1 by 162% compared to WT controls (**Figure 7D**), indicating that NL1 is bidirectionally modulated by MHCI.

**Figure 7.**
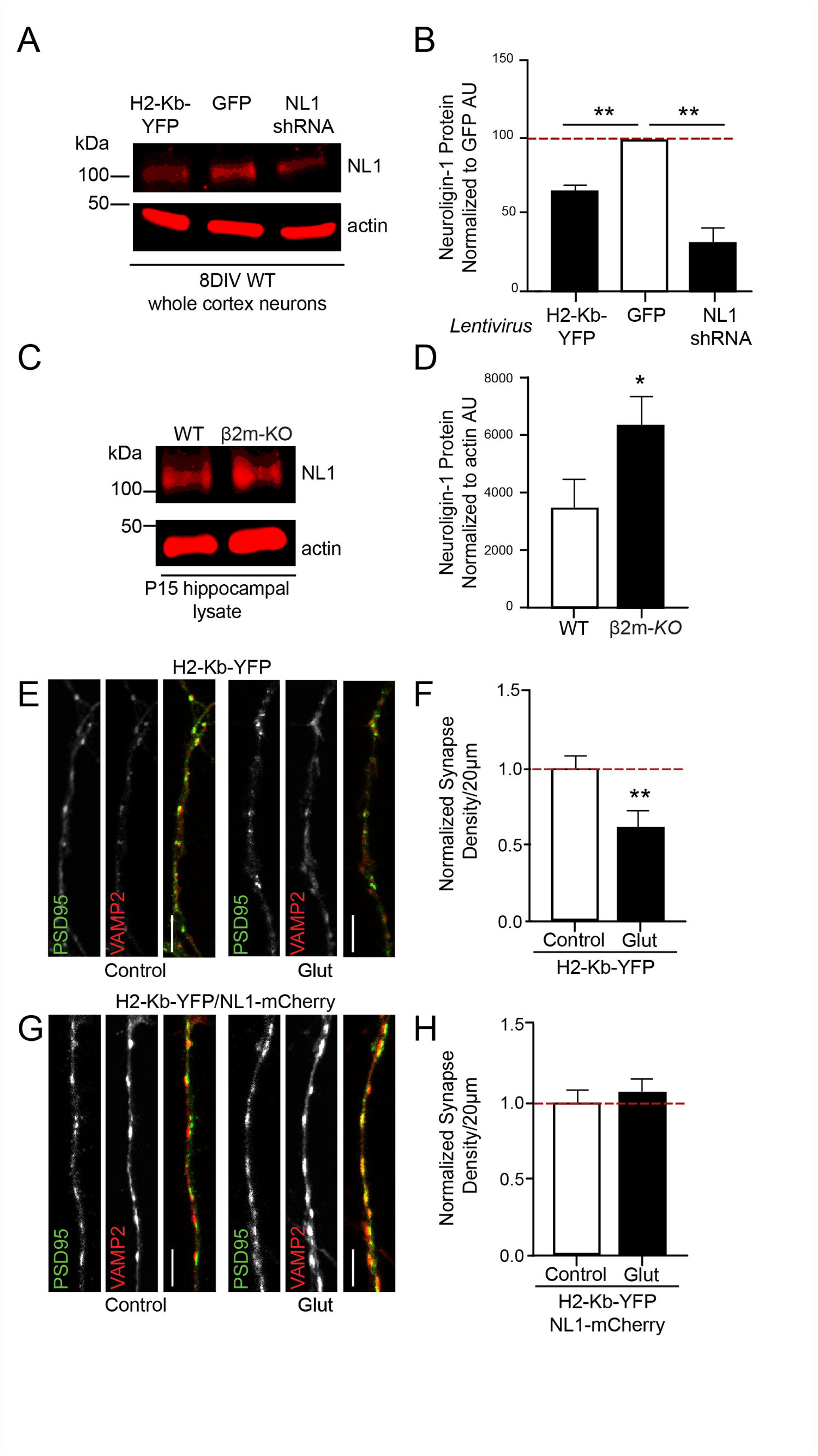
MHCI mediates glutamate-induced synapse loss by negatively regulating NL1 levels (A) Western blot of samples from lysed 8 div WT cortical neurons that were cultured and infected with lentivirus expressing GFP, H2-Kb-YFP, or NL1 shRNA at 3 div. (B) Protein levels normalized to actin from western blots as in (A) indicate that overexpression of H2-Kb-YFP significantly reduced NL1 total protein levels, as did NL1 shRNA. n = 4 cultures. **p<0.01, ANOVA. (C) Western blot of samples from lysed hippocampi from sex-matched littermate pairs of WT and β2m-KO mice. (D) Protein levels normalized to actin from western blots as in (C) indicate that loss of surface MHCI expression in the β2m-KO mice significantly increased NL1 total protein. N = 5 littermate pairs, *p<0.05, t-test. (E) Representative confocal images of WT occipital cortical mouse neurons transfected with H2-Kb-YFP that were incubated with glutamate for 10 min at 8 div, then fixed and stained with antibodies to postsynaptic PSD95 (*green*) and presynaptic VAMP2 puncta (*red*); glutamatergic synapses are defined as colocalized puncta. Scale bar, 5µm. (F) Glutamatergic synapses were significantly altered by glutamate treatment in the presence of excess H2-Kb-YFP. H2-Kb-YFP control N = 23, H2-Kb-YFP glut = 25 cells, **p<0.01, t-test. (G) Representative confocal images of WT occipital cortical mouse neurons transfected with H2-Kb-YFP and NL1-mCherry that were incubated with glutamate for 10 min at 8 div, then fixed and stained with antibodies to postsynaptic PSD95 (*green*) and presynaptic VAMP2 puncta (*red*); glutamatergic synapses are defined as colocalized puncta. Scale bar, 5µm. (D) Glutamatergic synapses were not altered by glutamate treatment in the presence of excess H2-Kb-YFP and excess NL1-mCherry. H2-Kb-YFP/NL1-mCherry control N = 27, H2-Kb-YFP glut = 27 cells, t-test.

Given that NL1 protein is negatively modulated by MHCI (**Figure 7A-D**) and NL1-Nrxn binding stabilizes synapses (**Figure 5D-E**), glutamate would be expected to cause synapse loss in neurons with elevated MHCI levels, phenocopying the effects of glutamate in NL1-KO neurons (**Figure 5G**). As predicted, there was a significant reduction in synapse density in WT occipital cortical mouse neurons that overexpressed H2-Kb-YFP after 10 min 50μM glutamate treatment (**Figure 7E-F**), reinforcing the necessity of NL1 in preventing glutamate-induced synapse loss.

Next, we investigated whether MHCI requires changes in NL1 levels to mediate the effects of glutamate. 3 div cortical neurons were transfected with both H2-Kb-YFP and NL1-mCherry before treatment with 10 min of glutamate or vehicle at 8 div and staining for PSD-95 and VAMP2 (**Figure 7G**). As predicted by our model, overexpression of NL1-mCherry together with H2-Kb-YFP phenocopied NL1-mCherry overexpression alone (**Figure 5A-B**) with no change in synapse density in response to glutamate (**Figure 7H**), indicating that MHCI overexpression affects glutamate-induced synapse loss through modulation of NL1 and suggesting that excess NL1 can stabilize a higher proportion of excitatory synapses even in the presence of glutamate and elevated MHCI.

## Discussion

Here, we describe a novel form of homeostatic plasticity that exists in young cortical neurons to rapidly alter glutamatergic synapse density in the face of changes in activity. Acute elevations in glutamate (10min) act as a dispersal signal for NMDARs and NL1 that reduces trafficking and causes loss of some synapses, selectively during an early period when connections are initially forming. This dispersal signal is opposed by trans-synaptic adhesion through binding of NL1 to Nrxn1, which stabilizes synapses. The effect of glutamate in causing synapse loss requires elevations in MHCI levels, which negatively regulate NL1 levels. In the presence of excess NL1 caused by either NL1 overexpression or loss of both primary forms of MHCI in C57BL/6 mice, or at sites of NL1 binding to presynaptic Nrxn1 in the hemi-synapse assay, glutamate does not cause synapse loss. In the absence (NL1-KO) or reduced presence of NL1 (MHCI overexpression), or at dendritic sites without binding of NL1 to Nrxn1, glutamate disperses NMDARs and causes glutamatergic synapse loss. Together, this glutamate-MHCI-NL1 signaling pathway represents a mechanism that could act to tightly regulate the number and location of the initial complement of synapses formed along developing dendrites.

Although homeostatic plasticity is essential for globally altering and resetting synaptic strength to counter the destabilizing effects of Hebbian plasticity in mature neural networks ^46,75,76^, the effects of acute global and local changes in neural activity on the initial establishment of neural networks has been unclear. Because synapses form in the absence of neurotransmitter release *in vivo* ^1-4,36^ and because synapse density is not altered by activity blockers in mature cultured hippocampal neurons ^30,46,75,77^, it has been assumed that neurotransmitter does not play a role in the initial formation of connections ^78^. In contrast, our results show that activation of NMDARs in young cortical networks regulates the density of glutamatergic synapses on a surprisingly short time-scale of minutes, but only in networks that are initially establishing their connections. The effects of glutamate were robust in young cortical neurons, leading to both a 35% decrease in glutamatergic synapse density and a 46% decrease in mEPSC frequency (with no change in mEPSC amplitude) after only 10min of glutamate treatment. Moreover, the glutamate-induced synapse loss occurred in both rat and mouse cultures, assessed by multiple combinations of colocalized proteins to define synapses (vGlut1/GluN2A/B and VAMP2/PSD-95). These changes occur on both a global and local level and are consistent with reports of high levels of spontaneous neurotransmitter release in developing tissue ^30-32^ as well as a lack of neurotransmitter transporters in young tissue ^79^, which could allow for greater levels of ambient neurotransmitter. The rapid and selective decrease in NRTP transport specifically at sites of glutamate perfusion suggests that local release of glutamate by axonal growth cones ^31-33^ could also regulate synapse formation. This new form of homeostatic plasticity resulting in rapid changes in synapse density during the initial formation of neural networks complements the emerging idea that networks tap into distinct forms of homeostatic plasticity at different stages of development throughout the lifespan ^80,81^. The rapid activity-induced changes in synapse density shown here may be necessary for neural networks to respond dynamically to rapidly changing input number as they connections initially form, while more established networks rely on altering synaptic weights or types of synaptic inputs to maintain their network stability over longer time-scales ^82^.

The central role for NMDARs in glutamate-induced synapse loss in young neural networks makes sense given that NMDARs are often the first glutamate receptors recruited to nascent synapses ^21^. In addition, the rapid trafficking of NRTPs within dendrites and their cycling with the membrane at pause sites before and during synapse formation ^27,42^ indicate that they are well-positioned to respond to local and global changes in glutamate. Indeed, both constitutive and agonist-induced internalization of NMDARs has been well characterized in heterologous cells and in young neural networks ^5,6,83-86^ and our data provide the first demonstration of a potential role for the glutamate released from axons before synapses form in NMDAR internalization and the initial establishment of connections ^30^. Our results do not directly address the GluN2 subunit specificity of this effect, but it is likely that GluN2B plays a central role given that GluN2B is the most abundant GluN2 subunit at young ages in cortex ^42,87-91^ and the extensive evidence that it undergoes rapid glutamate-induced internalization selectively in young neurons ^5,6^. Our results extend this work by showing that this glutamate-induced internalization of NMDARs occurs in conjunction with glutamatergic synapse loss and that it is accompanied by changes in the co-transport of NL1, a synaptic adhesion molecule that regulates NMDAR accumulation at synapses ^22,68,92^ and that has been shown to be regulated by activity ^93-96^. Given the likely central role of GluN2B in this process, it is possible that either the developmental subunit switch to primarily GluN2A-containing NMDARs ^91^, or the recruitment of synaptic scaffolding molecules like PSD-95 ^41^ may shift this novel form of homeostatic plasticity from altering synapse density in new neural networks to selectively altering synapse strength in more mature neural networks.

The role for glutamate in acting as a dispersal signal for NMDARs and a negative regulator of synapses in young networks is similar to the well-described role for ACh at the NMJ ^38,39^. At developing NMJs, ACh causes dispersal and internalization of AChRs unless an anti-dispersal signal, agrin, is present ^38,39^. Here, we show that glutamate causes internalization of NMDARs and synapse loss unless the receptors and synapses are stabilized by NL1. Similar to the role for ACh in destabilizing nascent ACh puncta in cultures and the role for glutamate activation of NMDARs at the NMJ causing synapse pruning during the initial stages of synaptogenesis ^40^, we found that glutamate causes internalization of NMDARs and glutamatergic synapse loss in young cortical cultures. Thus, the role for neurotransmitter in the formation of such disparate structures as the NMJ and cortical excitatory synapses appears to be conserved. Finally, it is likely that additional synaptic adhesion molecules play a similar role as NL1/Nrxn1 in opposing neurotransmitter-induced receptor internalization and synapse loss ^16,77,78^. If so, then this anti-dispersal function could contribute to the matching of presynaptic neurotransmitter with the accumulation of the correct type of neurotransmitter receptors at the synapse.

At first glance, our discovery that glutamate acts to cause synapse loss appears to be contradictory to the focus on glutamate (and neurotransmitters in general) in promoting spine and synapse formation ^10,11,37^. There are many studies showing that increasing activity causes dendritic spine formation in an NMDAR-dependent manner ^7-9,97,98^. Most relevant, the Sabatini lab showed that focal uncaging of glutamate ^10^ rapidly induces the formation of new dendritic spines with functional synapses selectively in young cortical slices. Importantly, NL1 is essential for this glutamate-induced synapse induction ^37^. Because these uncaging experiments were performed in intact tissue, which is tightly packed with neuronal processes, it is likely that any new dendritic spines stimulated to extend by local glutamate ^99-102^ will rapidly come into contact with presynaptic axons and thereby facilitate binding of NL1 to Nrxn1 to stabilize the nascent synapses and maintain them ^16,78^. In contrast, the low-density culture system used here limits the density of neighbouring axons, revealing that glutamate causes NMDAR internalization and synapse loss in the absence of presynaptic contact while mimicking presynaptic contacts with Nrxn1-expressing HEK cells prevents glutamate-induced synapse loss.

In addition to reconciling existing conflicting literature about the role for glutamate in the initial establishment of cortical connections, our experiments have revealed a novel molecular mechanism—neuronal MHCI—for glutamate in causing synapse loss. MHCI negatively regulates synapse density, selectively during the period of the initial establishment of cortical connections, which is the same developmental time window that glutamate rapidly alters synapse density ^43,45^. MHCI levels are known to be positively regulated by activity ^43,72,103,104^ and they contribute to synaptic scaling-like responses ^15,43,74,105^. In young cortical cultures, we previously showed that blocking activity with tetrodotoxin for two days decreases sMHCI and increases synapse density in an MHCI-dependent manner ^43^. Here, we show that sMHCI levels are rapidly increased to pre-existing sites by 10min glutamate in young cortical neurons, and that increase is required to decrease NL1 levels and limit synapse density. With few MHCI neuronal interactors known ^106-110^, this is the first demonstration that MHCI negatively regulates NL1 levels to control synapse density. Future studies will be needed to determine how MHCI alters NL1 and whether that interaction also plays a role in homeostatic plasticity of synaptic strength at later ages ^15,43,72,73,105,111^, and in synapse loss downstream of peripheral immune activation ^15,45,74,112-114^ and during neurodegeneration ^109,115^.

## Limitations of the study

While our discovery of the role for glutamate-MHCI-NL1-NMDARs in regulating early synapse loss in response to glutamate is significant, there are several limitations to be acknowledged. The primary potential limitation is that all of our experiments were performed using young primary neuronal cultures. This reduced approach was essential for determining the sufficiency of glutamate, at very early stages of network formation and in the absence of neighbouring presynaptic axons, in causing synapse loss. These experiments could not have been performed *in vivo*, but critically are consistent with and reconcile conflicting results from intact tissue. Future experiments will be needed to determine if all neurotransmitters act in a similar way to disperse neurotransmitter receptors during synapse formation. In addition, the source of glutamate and the role for astrocytes, which regulate neural activity and the initial stages of synapse formation ^116^, in this process remain open and important questions. Finally, novel live imaging tools will need to be developed to allow us to determine how rapidly and how locally, relative to the synapse, MHCI is altered by global and local changes in glutamate to cause synapse loss.

## Figure Legends

**Supplemental Figure 1.**
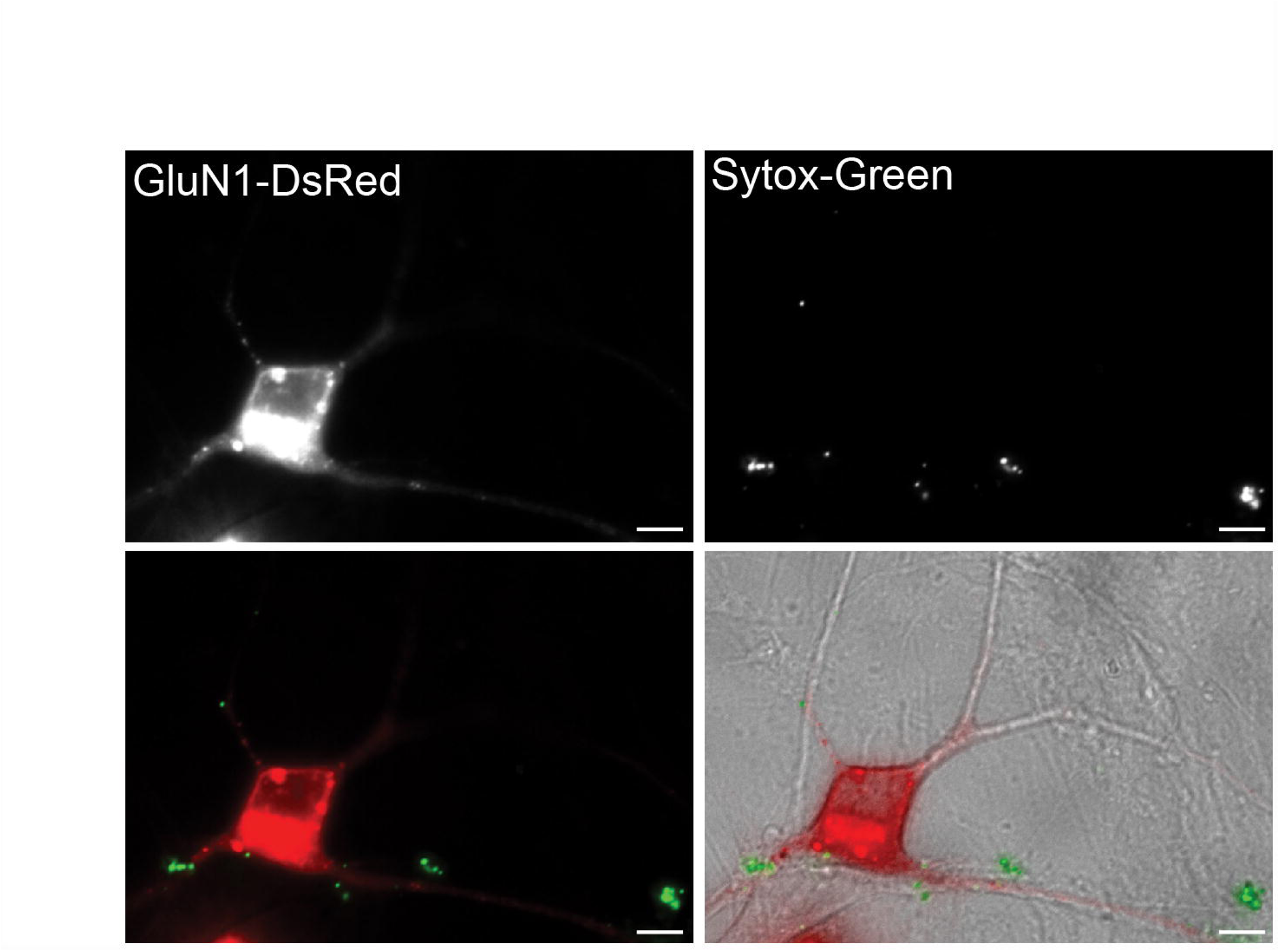
Acute glutamate treatment is not toxic to cultured neurons. Representative fluorescent images of 5 div cortical neurons expressing GluN1-DsRed (*left top*) and loaded with Sytox Green (0.5 µM) (*right top*) following incubation with glutamate (50 µM); overlay (*bottom left*). These images were overlaid on the brightfield image to show the morphology of neurons following glutamate treatment (*bottom right*). The representative cell shows a healthy soma and dendrites with typical GluN1-DsRed fluorescence and no detectable Sytox Green signal. Scale bar, 5µm.

**Supplemental Figure 2.**
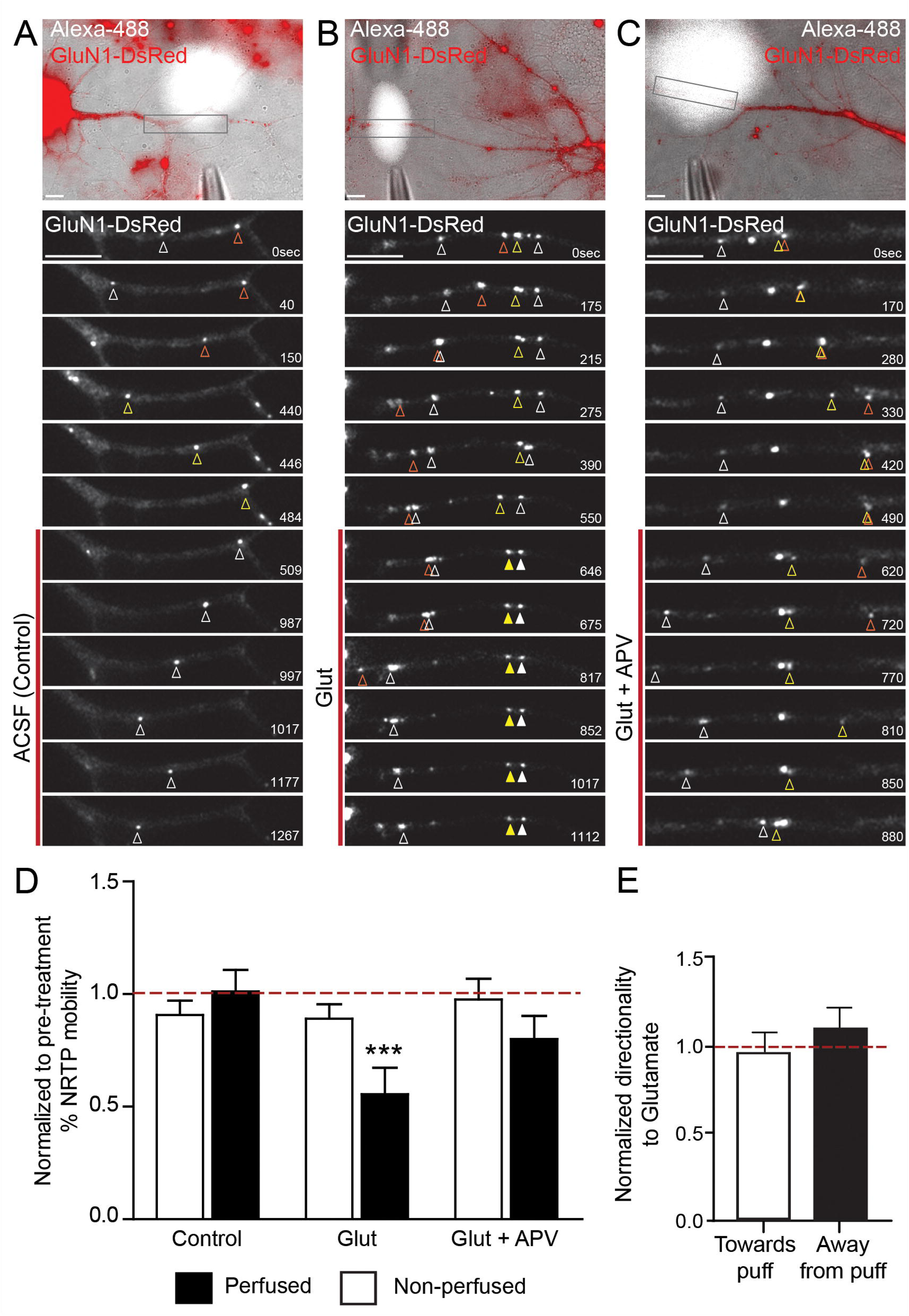
Glutamate acts locally to alter NRTP mobility in young cortical neurons. (A-C) Representative images showing a dendritic segment expressing GluN1-DsRed focally perfused with control ACSF (A), glutamate (B) or glutamate with APV (C). Five puffs of these solutions (2 sec duration, at 30 sec intervals) were perfused onto limited sections of dendrites for 10 min (pipette tip) using a picospritzer. Perfusion area is indicated by the Alexa-488 plume. Boxed regions at higher magnification below each image show changes in NRTP mobility before and during 10 min of puffing, indicated by the red line. Arrowheads indicate mobile NRTPs (*open arrowheads*) and stable NMDAR puncta (*closed arrowheads*) throughout the dendrite. Time in seconds. Scale bar, 5µm. (D) Focal perfusion of glutamate locally decreased NRTP mobility (*black bars*) compared to neighbouring, non-perfused dendritic segments (*open bars*). Perfusion of ACSF alone (control) had no effect on NRTP mobility but perfusion of APV prevented local inhibition of NRTP transport during focal perfusion of glutamate. Control (ACSF) n = 5, Glut = 6, Glut + APV = 5. Data is plotted as mean +/- SEM, normalized to GluN1 mobility within dendrite before drug treatment (*control black bar; red dotted line*). ***p<0.001, ANOVA. (E) Focal perfusion of glutamate does not attract NRTPs towards or away from the focal stream in neighbouring, non-perfused dendritic segments. Data is plotted as mean +/- SEM, normalized to control mobility prior to focal perfusion (*red dotted line*). n = 6 cells, t-test.

**Supplemental Figure 3.**
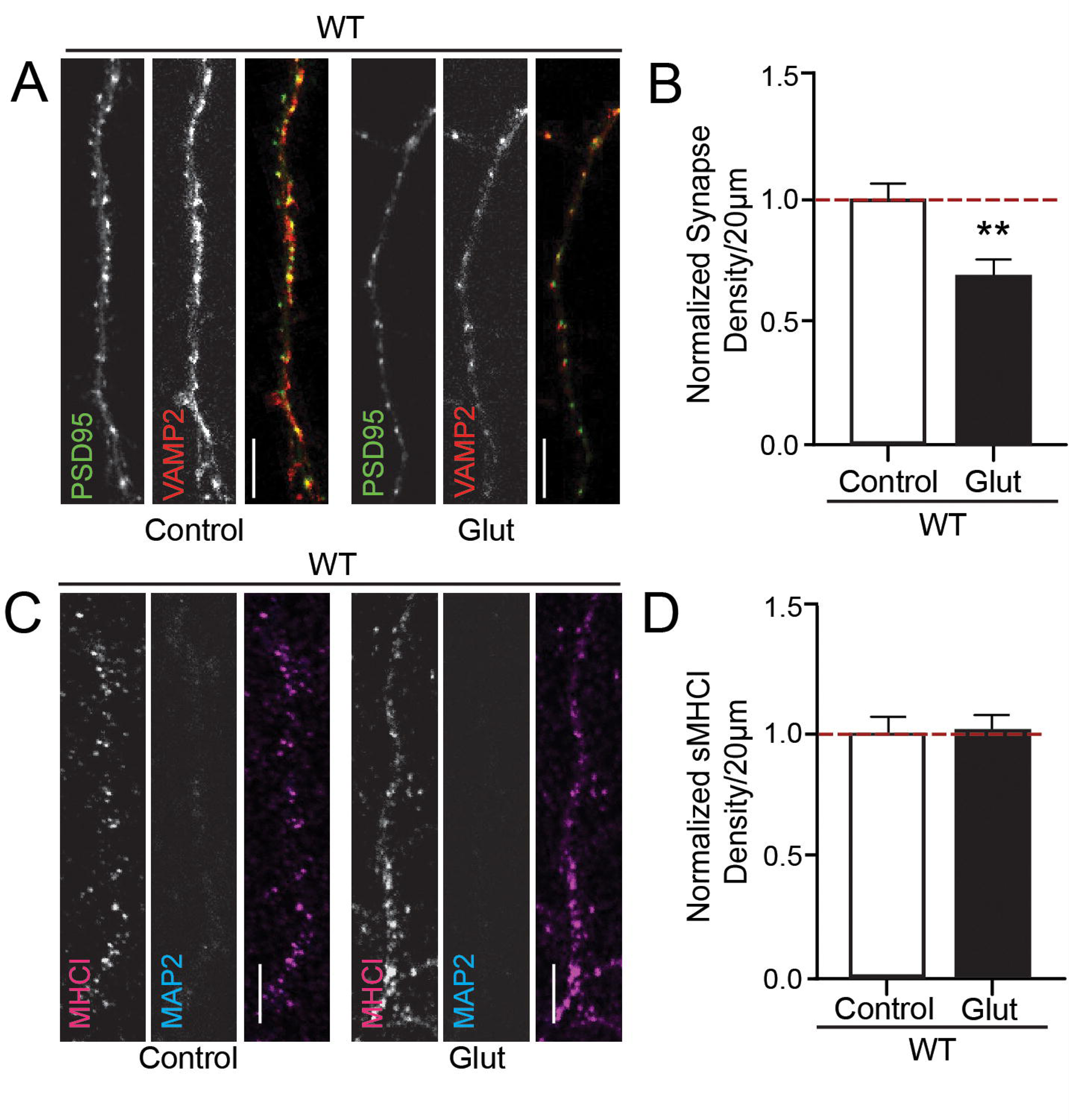
Acute glutamate treatment causes synapse loss in young mouse cultures but does not alter density of sMHCI punta. (A) Representative confocal images of WT occipital cortical mouse neurons at 8 div. Neurons were fixed and stained with antibodies to identify glutamatergic synapses defined as colocalized puncta of postsynaptic PSD-95 (*green*) and presynaptic VAMP2 (*red*). Scale bar, 5µm. (B) Treatment with 10 minutes of glutamate significantly reduced synapse density in WT murine neurons. Control n = 34 cells, glutamate = 23 cells. Data is plotted as mean +/- SEM, normalized to control cells from the same culture/experiment (*red dotted line*). ** p < 0.01, t-test. (C) Representative confocal images of non-permeabilized WT occipital cortical mouse neurons at 8 div. Neurons were fixed and stained with antibodies to identify sMHCI (*magenta*) in the absence of permeabilization as defined as no MAP2 (*blue)* colocalization. Scale bar, 5µm. (D) There was no significant change in sMHCI punctal density in response to glutamate. Control n = 21 cells, glutamate n = 27 cells, Data is plotted as mean +/- SEM, normalized to control cells from the same culture/experiment (*red dotted line*). T-test.

## References

1. Sigler, A., Oh, W.C., Imig, C., Altas, B., Kawabe, H., Cooper, B.H., Kwon, H.B., Rhee, J.S., and Brose, N. (2017). Formation and Maintenance of Functional Spines in the Absence of Presynaptic Glutamate Release. Neuron 94, 304–311 e304. 10.1016/j.neuron.2017.03.029.

2. Sando, R., Bushong, E., Zhu, Y., Huang, M., Considine, C., Phan, S., Ju, S., Uytiepo, M., Ellisman, M., and Maximov, A. (2017). Assembly of Excitatory Synapses in the Absence of Glutamatergic Neurotransmission. Neuron 94, 312–321 e313. 10.1016/j.neuron.2017.03.047.

3. Verhage, M., Maia, A.S., Plomp, J.J., Brussaard, A.B., Heeroma, J.H., Vermeer, H., Toonen, R.F., Hammer, R.E., van den Berg, T.K., Missler, M., et al. (2000). Synaptic assembly of the brain in the absence of neurotransmitter secretion. Science 287, 864–869. 10.1126/science.287.5454.864.

4. Held, R.G., Liu, C., Ma, K., Ramsey, A.M., Tarr, T.B., De Nola, G., Wang, S.S.H., Wang, J., van den Maagdenberg, A., Schneider, T., et al. (2020). Synapse and Active Zone Assembly in the Absence of Presynaptic Ca(2+) Channels and Ca(2+) Entry. Neuron 107, 667–683 e669. 10.1016/j.neuron.2020.05.032.

5. Nong, Y., Huang, Y.Q., and Salter, M.W. (2004). NMDA receptors are movin’ in. Current opinion in neurobiology 14, 353–361. 10.1016/j.conb.2004.05.001.

6. Roche, K.W., Standley, S., McCallum, J., Dune Ly, C., Ehlers, M.D., and Wenthold, R.J. (2001). Molecular determinants of NMDA receptor internalization. Nature neuroscience 4, 794–802. 10.1038/90498.

7. Engert, F., and Bonhoeffer, T. (1999). Dendritic spine changes associated with hippocampal long-term synaptic plasticity. Nature 399, 66–70. 10.1038/19978.

8. Maletic-Savatic, M., Malinow, R., and Svoboda, K. (1999). Rapid dendritic morphogenesis in CA1 hippocampal dendrites induced by synaptic activity. Science 283, 1923–1927. 10.1126/science.283.5409.1923.

9. Toni, N., Buchs, P.A., Nikonenko, I., Bron, C.R., and Muller, D. (1999). LTP promotes formation of multiple spine synapses between a single axon terminal and a dendrite. Nature 402, 421–425. 10.1038/46574.

10. Kwon, H.B., and Sabatini, B.L. (2011). Glutamate induces de novo growth of functional spines in developing cortex. Nature 474, 100–104. 10.1038/nature09986.

11. Oh, W.C., Lutzu, S., Castillo, P.E., and Kwon, H.B. (2016). De novo synaptogenesis induced by GABA in the developing mouse cortex. Science 353, 1037–1040. 10.1126/science.aaf5206.

12. Grant, S.G. (2012). Synaptopathies: diseases of the synaptome. Current opinion in neurobiology 22, 522–529. 10.1016/j.conb.2012.02.002.

13. Melom, J.E., and Littleton, J.T. (2011). Synapse development in health and disease. Current opinion in genetics & development 21, 256–261. 10.1016/j.gde.2011.01.002.

14. Washbourne, P. (2015). Synapse assembly and neurodevelopmental disorders. Neuropsychopharmacology 40, 4–15. 10.1038/npp.2014.163.

15. McAllister, A.K. (2014). Major histocompatibility complex I in brain development and schizophrenia. Biol Psychiatry 75, 262–268. 10.1016/j.biopsych.2013.10.003.

16. Sudhof, T.C. (2018). Towards an Understanding of Synapse Formation. Neuron 100, 276–293. 10.1016/j.neuron.2018.09.040.

17. Cameron, S., and McAllister, A.K. (2018). Immunoglobulin-Like Receptors and Their Impact on Wiring of Brain Synapses. Annu Rev Genet 52, 567–590. 10.1146/annurev-genet-120417-031513.

18. McAllister, A.K. (2007). Dynamic aspects of CNS synapse formation. Annu Rev Neurosci 30, 425–450. 10.1146/annurev.neuro.29.051605.112830.

19. Waites, C.L., Craig, A.M., and Garner, C.C. (2005). Mechanisms of vertebrate synaptogenesis. Annu Rev Neurosci 28, 251–274. 10.1146/annurev.neuro.27.070203.144336.

20. Tallafuss, A., Constable, J.R., and Washbourne, P. (2010). Organization of central synapses by adhesion molecules. Eur J Neurosci 32, 198–206. 10.1111/j.1460-9568.2010.07340.x.

21. Washbourne, P., Bennett, J.E., and McAllister, A.K. (2002). Rapid recruitment of NMDA receptor transport packets to nascent synapses. Nature neuroscience 5, 751–759. 10.1038/nn883.

22. Barrow, S.L., Constable, J.R., Clark, E., El-Sabeawy, F., McAllister, A.K., and Washbourne, P. (2009). Neuroligin1: a cell adhesion molecule that recruits PSD-95 and NMDA receptors by distinct mechanisms during synaptogenesis. Neural development 4, 17. 10.1186/1749-8104-4-17.

23. Wu, X., Morishita, W.K., Riley, A.M., Hale, W.D., Sudhof, T.C., and Malenka, R.C. (2019). Neuroligin-1 Signaling Controls LTP and NMDA Receptors by Distinct Molecular Pathways. Neuron 102, 621–635 e623. 10.1016/j.neuron.2019.02.013.

24. Wei, Z., Behrman, B., Wu, W.H., and Chen, B.S. (2015). Subunit-specific regulation of N-methyl-D-aspartate (NMDA) receptor trafficking by SAP102 protein splice variants. J Biol Chem 290, 5105–5116. 10.1074/jbc.M114.599969.

25. Behuet, S., Cremer, J.N., Cremer, M., Palomero-Gallagher, N., Zilles, K., and Amunts, K. (2019). Developmental Changes of Glutamate and GABA Receptor Densities in Wistar Rats. Front Neuroanat 13, 100. 10.3389/fnana.2019.00100.

26. Groc, L., Choquet, D., Stephenson, F.A., Verrier, D., Manzoni, O.J., and Chavis, P. (2007). NMDA receptor surface trafficking and synaptic subunit composition are developmentally regulated by the extracellular matrix protein Reelin. J Neurosci 27, 10165–10175. 10.1523/JNEUROSCI.1772-07.2007.

27. Washbourne, P., Liu, X.B., Jones, E.G., and McAllister, A.K. (2004). Cycling of NMDA receptors during trafficking in neurons before synapse formation. J Neurosci 24, 8253–8264. 10.1523/JNEUROSCI.2555-04.2004.

28. Hume, R.I., Role, L.W., and Fischbach, G.D. (1983). Acetylcholine release from growth cones detected with patches of acetylcholine receptor-rich membranes. Nature 305, 632–634. 10.1038/305632a0.

29. Young, S.H., and Poo, M.M. (1983). Spontaneous release of transmitter from growth cones of embryonic neurones. Nature 305, 634–637. 10.1038/305634a0.

30. Andreae, L.C., and Burrone, J. (2018). The role of spontaneous neurotransmission in synapse and circuit development. J Neurosci Res 96, 354–359. 10.1002/jnr.24154.

31. Matteoli, M., Takei, K., Perin, M.S., Sudhof, T.C., and De Camilli, P. (1992). Exo-endocytotic recycling of synaptic vesicles in developing processes of cultured hippocampal neurons. J Cell Biol 117, 849–861. 10.1083/jcb.117.4.849.

32. Sabo, S.L., and McAllister, A.K. (2003). Mobility and cycling of synaptic protein-containing vesicles in axonal growth cone filopodia. Nature neuroscience 6, 1264–1269. 10.1038/nn1149.

33. Sabo, S.L., Gomes, R.A., and McAllister, A.K. (2006). Formation of presynaptic terminals at predefined sites along axons. J Neurosci 26, 10813–10825. 10.1523/JNEUROSCI.2052-06.2006.

34. Katz, L.C., and Crowley, J.C. (2002). Development of cortical circuits: lessons from ocular dominance columns. Nat Rev Neurosci 3, 34–42. 10.1038/nrn703.

35. Hooks, B.M., and Chen, C. (2020). Circuitry Underlying Experience-Dependent Plasticity in the Mouse Visual System. Neuron 107, 986–987. 10.1016/j.neuron.2020.08.004.

36. Varoqueaux, F., Sigler, A., Rhee, J.S., Brose, N., Enk, C., Reim, K., and Rosenmund, C. (2002). Total arrest of spontaneous and evoked synaptic transmission but normal synaptogenesis in the absence of Munc13-mediated vesicle priming. Proceedings of the National Academy of Sciences of the United States of America 99, 9037–9042. 10.1073/pnas.122623799.

37. Kwon, H.B., Kozorovitskiy, Y., Oh, W.J., Peixoto, R.T., Akhtar, N., Saulnier, J.L., Gu, C., and Sabatini, B.L. (2012). Neuroligin-1-dependent competition regulates cortical synaptogenesis and synapse number. Nature neuroscience 15, 1667–1674. 10.1038/nn.3256.

38. Misgeld, T., Kummer, T.T., Lichtman, J.W., and Sanes, J.R. (2005). Agrin promotes synaptic differentiation by counteracting an inhibitory effect of neurotransmitter. Proceedings of the National Academy of Sciences of the United States of America 102, 11088–11093. 10.1073/pnas.0504806102.

39. Rodriguez Cruz, P.M., Cossins, J., Beeson, D., and Vincent, A. (2020). The Neuromuscular Junction in Health and Disease: Molecular Mechanisms Governing Synaptic Formation and Homeostasis. Front Mol Neurosci 13, 610964. 10.3389/fnmol.2020.610964.

40. Personius, K.E., Slusher, B.S., and Udin, S.B. (2016). Neuromuscular NMDA Receptors Modulate Developmental Synapse Elimination. J Neurosci 36, 8783–8789. 10.1523/JNEUROSCI.1181-16.2016.

41. Lavezzari, G., McCallum, J., Dewey, C.M., and Roche, K.W. (2004). Subunit-specific regulation of NMDA receptor endocytosis. J Neurosci 24, 6383–6391. 10.1523/JNEUROSCI.1890-04.2004.

42. Tovar, K.R., and Westbrook, G.L. (2002). Mobile NMDA receptors at hippocampal synapses. Neuron 34, 255–264. 10.1016/s0896-6273(02)00658-x.

43. Glynn, M.W., Elmer, B.M., Garay, P.A., Liu, X.B., Needleman, L.A., El-Sabeawy, F., and McAllister, A.K. (2011). MHCI negatively regulates synapse density during the establishment of cortical connections. Nature neuroscience 14, 442–451. 10.1038/nn.2764.

44. Needleman, L.A., Liu, X.B., El-Sabeawy, F., Jones, E.G., and McAllister, A.K. (2010). MHC class I molecules are present both pre- and postsynaptically in the visual cortex during postnatal development and in adulthood. Proceedings of the National Academy of Sciences of the United States of America 107, 16999–17004. 10.1073/pnas.1006087107.

45. Elmer, B.M., Estes, M.L., Barrow, S.L., and McAllister, A.K. (2013). MHCI requires MEF2 transcription factors to negatively regulate synapse density during development and in disease. J Neurosci 33, 13791–13804. 10.1523/JNEUROSCI.2366-13.2013.

46. Turrigiano, G. (2012). Homeostatic synaptic plasticity: local and global mechanisms for stabilizing neuronal function. Cold Spring Harbor perspectives in biology 4, a005736. 10.1101/cshperspect.a005736.

47. Blundell, J., Blaiss, C.A., Etherton, M.R., Espinosa, F., Tabuchi, K., Walz, C., Bolliger, M.F., Sudhof, T.C., and Powell, C.M. (2010). Neuroligin-1 deletion results in impaired spatial memory and increased repetitive behavior. J Neurosci 30, 2115–2129. 10.1523/JNEUROSCI.4517-09.2010.

48. Boulanger, L.M., and Shatz, C.J. (2004). Immune signalling in neural development, synaptic plasticity and disease. Nat Rev Neurosci 5, 521–531. 10.1038/nrn1428.

49. Vugmeyster, Y., Glas, R., Perarnau, B., Lemonnier, F.A., Eisen, H., and Ploegh, H. (1998). Major histocompatibility complex (MHC) class I KbDb -/- deficient mice possess functional CD8+ T cells and natural killer cells. Proceedings of the National Academy of Sciences of the United States of America 95, 12492–12497. 10.1073/pnas.95.21.12492.

50. Ko, J., Fuccillo, M.V., Malenka, R.C., and Sudhof, T.C. (2009). LRRTM2 functions as a neurexin ligand in promoting excitatory synapse formation. Neuron 64, 791–798. 10.1016/j.neuron.2009.12.012.

51. Takahashi, H., Katayama, K., Sohya, K., Miyamoto, H., Prasad, T., Matsumoto, Y., Ota, M., Yasuda, H., Tsumoto, T., Aruga, J., and Craig, A.M. (2012). Selective control of inhibitory synapse development by Slitrk3-PTPdelta trans-synaptic interaction. Nature neuroscience 15, 389–398, S381-382. 10.1038/nn.3040.

52. Kopec, C.D., Li, B., Wei, W., Boehm, J., and Malinow, R. (2006). Glutamate receptor exocytosis and spine enlargement during chemically induced long-term potentiation. J Neurosci 26, 2000–2009. 10.1523/JNEUROSCI.3918-05.2006.

53. Fu, Z., Washbourne, P., Ortinski, P., and Vicini, S. (2003). Functional excitatory synapses in HEK293 cells expressing neuroligin and glutamate receptors. J Neurophysiol 90, 3950–3957. 10.1152/jn.00647.2003.

54. Desch, K., Schuman, E.M., and Langer, J.D. (2022). Quantifying phosphorylation dynamics in primary neuronal cultures using LC-MS/MS. STAR Protoc 3, 101063. 10.1016/j.xpro.2021.101063.

55. Friedman, H.V., Bresler, T., Garner, C.C., and Ziv, N.E. (2000). Assembly of new individual excitatory synapses: time course and temporal order of synaptic molecule recruitment. Neuron 27, 57–69.

56. Rao, A., Kim, E., Sheng, M., and Craig, A.M. (1998). Heterogeneity in the molecular composition of excitatory postsynaptic sites during development of hippocampal neurons in culture. J Neurosci 18, 1217–1229. 10.1523/JNEUROSCI.18-04-01217.1998.

57. Okawa, H., Hoon, M., Yoshimatsu, T., Della Santina, L., and Wong, R.O.L. (2014). Illuminating the multifaceted roles of neurotransmission in shaping neuronal circuitry. Neuron 83, 1303–1318. 10.1016/j.neuron.2014.08.029.

58. Evans, G.J., and Cousin, M.A. (2007). Simultaneous monitoring of three key neuronal functions in primary neuronal cultures. Journal of neuroscience methods 160, 197–205. 10.1016/j.jneumeth.2006.09.012.

59. Ashby, M.C., Ibaraki, K., and Henley, J.M. (2004). It’s green outside: tracking cell surface proteins with pH-sensitive GFP. Trends in neurosciences 27, 257–261. 10.1016/j.tins.2004.03.010.

60. Zucker, R.S. (1999). Calcium- and activity-dependent synaptic plasticity. Current opinion in neurobiology 9, 305–313.

61. Spitzer, N.C. (2002). Activity-dependent neuronal differentiation prior to synapse formation: the functions of calcium transients. Journal of physiology, Paris 96, 73–80.

62. Connor, J.A., Petrozzino, J., Pozzo-Miller, L.D., and Otani, S. (1999). Calcium signals in long-term potentiation and long-term depression. Canadian journal of physiology and pharmacology 77, 722–734.

63. Cummings, J.A., Mulkey, R.M., Nicoll, R.A., and Malenka, R.C. (1996). Ca2+ signaling requirements for long-term depression in the hippocampus. Neuron 16, 825–833.

64. Budreck, E.C., Kwon, O.B., Jung, J.H., Baudouin, S., Thommen, A., Kim, H.S., Fukazawa, Y., Harada, H., Tabuchi, K., Shigemoto, R., et al. (2013). Neuroligin-1 controls synaptic abundance of NMDA-type glutamate receptors through extracellular coupling. Proceedings of the National Academy of Sciences of the United States of America 110, 725–730. 10.1073/pnas.1214718110.

65. Varoqueaux, F., Aramuni, G., Rawson, R.L., Mohrmann, R., Missler, M., Gottmann, K., Zhang, W., Sudhof, T.C., and Brose, N. (2006). Neuroligins determine synapse maturation and function. Neuron 51, 741–754. 10.1016/j.neuron.2006.09.003.

66. Chen, L.Y., Jiang, M., Zhang, B., Gokce, O., and Sudhof, T.C. (2017). Conditional Deletion of All Neurexins Defines Diversity of Essential Synaptic Organizer Functions for Neurexins. Neuron 94, 611–625 e614. 10.1016/j.neuron.2017.04.011.

67. Tsetsenis, T., Boucard, A.A., Arac, D., Brunger, A.T., and Sudhof, T.C. (2014). Direct visualization of trans-synaptic neurexin-neuroligin interactions during synapse formation. J Neurosci 34, 15083–15096. 10.1523/JNEUROSCI.0348-14.2014.

68. Chih, B., Engelman, H., and Scheiffele, P. (2005). Control of excitatory and inhibitory synapse formation by neuroligins. Science 307, 1324–1328. 10.1126/science.1107470.

69. Scheiffele, P., Fan, J.H., Choih, J., Fetter, R., and Serafini, T. (2000). Neuroligin expressed in nonneuronal cells triggers presynaptic development in contacting axons. Cell 101, 657–669. Doi 10.1016/S0092-8674(00)80877-6.

70. Biederer, T., and Scheiffele, P. (2007). Mixed-culture assays for analyzing neuronal synapse formation. Nature protocols 2, 670–676. 10.1038/nprot.2007.92.

71. Graf, E.R., Zhang, X., Jin, S.X., Linhoff, M.W., and Craig, A.M. (2004). Neurexins induce differentiation of GABA and glutamate postsynaptic specializations via neuroligins. Cell 119, 1013–1026. 10.1016/j.cell.2004.11.035.

72. Corriveau, R.A., Huh, G.S., and Shatz, C.J. (1998). Regulation of class I MHC gene expression in the developing and mature CNS by neural activity. Neuron 21, 505–520.

73. Huh, G.S., Boulanger, L.M., Du, H., Riquelme, P.A., Brotz, T.M., and Shatz, C.J. (2000). Functional requirement for class I MHC in CNS development and plasticity. Science 290, 2155–2159.

74. Elmer, B.M., and McAllister, A.K. (2012). Major histocompatibility complex class I proteins in brain development and plasticity. Trends in neurosciences 35, 660–670. 10.1016/j.tins.2012.08.001.

75. Turrigiano, G.G. (2017). The dialectic of Hebb and homeostasis. Philos Trans R Soc Lond B Biol Sci 372. 10.1098/rstb.2016.0258.

76. Chen, L., Li, X., Tjia, M., and Thapliyal, S. (2022). Homeostatic plasticity and excitation-inhibition balance: The good, the bad, and the ugly. Current opinion in neurobiology 75, 102553. 10.1016/j.conb.2022.102553.

77. Ko, J., Soler-Llavina, G.J., Fuccillo, M.V., Malenka, R.C., and Sudhof, T.C. (2011). Neuroligins/LRRTMs prevent activity- and Ca2+/calmodulin-dependent synapse elimination in cultured neurons. J Cell Biol 194, 323–334. 10.1083/jcb.201101072.

78. Sudhof, T.C. (2021). The cell biology of synapse formation. J Cell Biol 220. 10.1083/jcb.202103052.

79. Thomas, C.G., Tian, H., and Diamond, J.S. (2011). The relative roles of diffusion and uptake in clearing synaptically released glutamate change during early postnatal development. J Neurosci 31, 4743–4754. 10.1523/JNEUROSCI.5953-10.2011.

80. Wierenga, C.J., Walsh, M.F., and Turrigiano, G.G. (2006). Temporal regulation of the expression locus of homeostatic plasticity. J Neurophysiol 96, 2127–2133. 10.1152/jn.00107.2006.

81. Wen, W., and Turrigiano, G.G. (2021). Developmental Regulation of Homeostatic Plasticity in Mouse Primary Visual Cortex. J Neurosci 41, 9891–9905. 10.1523/JNEUROSCI.1200-21.2021.

82. Turrigiano, G. (2011). Too many cooks? Intrinsic and synaptic homeostatic mechanisms in cortical circuit refinement. Annu Rev Neurosci 34, 89–103. 10.1146/annurev-neuro-060909-153238.

83. Carroll, R.C., Beattie, E.C., Xia, H., Luscher, C., Altschuler, Y., Nicoll, R.A., Malenka, R.C., and von Zastrow, M. (1999). Dynamin-dependent endocytosis of ionotropic glutamate receptors. Proceedings of the National Academy of Sciences of the United States of America 96, 14112–14117. 10.1073/pnas.96.24.14112.

84. Nong, Y., Huang, Y.Q., Ju, W., Kalia, L.V., Ahmadian, G., Wang, Y.T., and Salter, M.W. (2003). Glycine binding primes NMDA receptor internalization. Nature 422, 302–307. 10.1038/nature01497.

85. Sans, N., Prybylowski, K., Petralia, R.S., Chang, K., Wang, Y.X., Racca, C., Vicini, S., and Wenthold, R.J. (2003). NMDA receptor trafficking through an interaction between PDZ proteins and the exocyst complex. Nat Cell Biol 5, 520–530. 10.1038/ncb990.

86. Vissel, B., Krupp, J.J., Heinemann, S.F., and Westbrook, G.L. (2001). A use-dependent tyrosine dephosphorylation of NMDA receptors is independent of ion flux. Nature neuroscience 4, 587–596. 10.1038/88404.

87. Monyer, H., Burnashev, N., Laurie, D.J., Sakmann, B., and Seeburg, P.H. (1994). Developmental and regional expression in the rat brain and functional properties of four NMDA receptors. Neuron 12, 529–540. 10.1016/0896-6273(94)90210-0.

88. Rumbaugh, G., and Vicini, S. (1999). Distinct synaptic and extrasynaptic NMDA receptors in developing cerebellar granule neurons. J Neurosci 19, 10603–10610. 10.1523/JNEUROSCI.19-24-10603.1999.

89. Sheng, M., Cummings, J., Roldan, L.A., Jan, Y.N., and Jan, L.Y. (1994). Changing subunit composition of heteromeric NMDA receptors during development of rat cortex. Nature 368, 144–147. 10.1038/368144a0.

90. Vicini, S., Wang, J.F., Li, J.H., Zhu, W.J., Wang, Y.H., Luo, J.H., Wolfe, B.B., and Grayson, D.R. (1998). Functional and pharmacological differences between recombinant N-methyl-D-aspartate receptors. J Neurophysiol 79, 555–566. 10.1152/jn.1998.79.2.555.

91. Barria, A., and Malinow, R. (2002). Subunit-specific NMDA receptor trafficking to synapses. Neuron 35, 345–353. 10.1016/s0896-6273(02)00776-6.

92. Chubykin, A.A., Atasoy, D., Etherton, M.R., Brose, N., Kavalali, E.T., Gibson, J.R., and Sudhof, T.C. (2007). Activity-dependent validation of excitatory versus inhibitory synapses by neuroligin-1 versus neuroligin-2. Neuron 54, 919–931. 10.1016/j.neuron.2007.05.029.

93. Bemben, M.A., Shipman, S.L., Hirai, T., Herring, B.E., Li, Y., Badger, J.D., 2nd, Nicoll, R.A., Diamond, J.S., and Roche, K.W. (2014). CaMKII phosphorylation of neuroligin-1 regulates excitatory synapses. Nature neuroscience 17, 56-64. 10.1038/nn.3601.

94. Peixoto, R.T., Kunz, P.A., Kwon, H., Mabb, A.M., Sabatini, B.L., Philpot, B.D., and Ehlers, M.D. (2012). Transsynaptic signaling by activity-dependent cleavage of neuroligin-1. Neuron 76, 396–409. 10.1016/j.neuron.2012.07.006.

95. Schapitz, I.U., Behrend, B., Pechmann, Y., Lappe-Siefke, C., Kneussel, S.J., Wallace, K.E., Stempel, A.V., Buck, F., Grant, S.G., Schweizer, M., et al. (2010). Neuroligin 1 is dynamically exchanged at postsynaptic sites. J Neurosci 30, 12733–12744. 10.1523/JNEUROSCI.0896-10.2010.

96. Suzuki, K., Hayashi, Y., Nakahara, S., Kumazaki, H., Prox, J., Horiuchi, K., Zeng, M., Tanimura, S., Nishiyama, Y., Osawa, S., et al. (2012). Activity-dependent proteolytic cleavage of neuroligin-1. Neuron 76, 410–422. 10.1016/j.neuron.2012.10.003.

97. Nagerl, U.V., Eberhorn, N., Cambridge, S.B., and Bonhoeffer, T. (2004). Bidirectional activity-dependent morphological plasticity in hippocampal neurons. Neuron 44, 759–767. 10.1016/j.neuron.2004.11.016.

98. Jourdain, P., Fukunaga, K., and Muller, D. (2003). Calcium/calmodulin-dependent protein kinase II contributes to activity-dependent filopodia growth and spine formation. J Neurosci 23, 10645–10649. 10.1523/JNEUROSCI.23-33-10645.2003.

99. Cruz-Martin, A., Crespo, M., and Portera-Cailliau, C. (2012). Glutamate induces the elongation of early dendritic protrusions via mGluRs in wild type mice, but not in fragile X mice. PLoS One 7, e32446. 10.1371/journal.pone.0032446.

100. Dailey, M.E., and Smith, S.J. (1996). The dynamics of dendritic structure in developing hippocampal slices. J Neurosci 16, 2983–2994. 10.1523/JNEUROSCI.16-09-02983.1996.

101. Portera-Cailliau, C., Pan, D.T., and Yuste, R. (2003). Activity-regulated dynamic behavior of early dendritic protrusions: evidence for different types of dendritic filopodia. J Neurosci 23, 7129–7142. 10.1523/JNEUROSCI.23-18-07129.2003.

102. Wong, W.T., and Wong, R.O. (2001). Changing specificity of neurotransmitter regulation of rapid dendritic remodeling during synaptogenesis. Nature neuroscience 4, 351–352. 10.1038/85987.

103. Lv, D., Shen, Y., Peng, Y., Liu, J., Miao, F., and Zhang, J. (2015). Neuronal MHC Class I Expression Is Regulated by Activity Driven Calcium Signaling. PLoS One 10, e0135223. 10.1371/journal.pone.0135223.

104. Ribic, A., Flugge, G., Schlumbohm, C., Matz-Rensing, K., Walter, L., and Fuchs, E. (2011). Activity-dependent regulation of MHC class I expression in the developing primary visual cortex of the common marmoset monkey. Behav Brain Funct 7, 1. 10.1186/1744-9081-7-1.

105. Goddard, C.A., Butts, D.A., and Shatz, C.J. (2007). Regulation of CNS synapses by neuronal MHC class I. Proceedings of the National Academy of Sciences of the United States of America 104, 6828–6833. 10.1073/pnas.0702023104.

106. Kim, T., Vidal, G.S., Djurisic, M., William, C.M., Birnbaum, M.E., Garcia, K.C., Hyman, B.T., and Shatz, C.J. (2013). Human LilrB2 is a beta-amyloid receptor and its murine homolog PirB regulates synaptic plasticity in an Alzheimer’s model. Science 341, 1399–1404. 10.1126/science.1242077.

107. Dixon-Salazar, T.J., Fourgeaud, L., Tyler, C.M., Poole, J.R., Park, J.J., and Boulanger, L.M. (2014). MHC class I limits hippocampal synapse density by inhibiting neuronal insulin receptor signaling. J Neurosci 34, 11844–11856. 10.1523/JNEUROSCI.4642-12.2014.

108. Frietze, K.K., Pappy, A.L., 2nd, Melson, J.W., O’Driscoll, E.E., Tyler, C.M., Perlman, D.H., and Boulanger, L.M. (2016). Cryptic protein-protein interaction motifs in the cytoplasmic domain of MHCI proteins. BMC Immunol 17, 24. 10.1186/s12865-016-0154-z.

109. Kim, M.S., Cho, K., Cho, M.H., Kim, N.Y., Kim, K., Kim, D.H., and Yoon, S.Y. (2023). Neuronal MHC-I complex is destabilized by amyloid-beta and its implications in Alzheimer’s disease. Cell Biosci 13, 181. 10.1186/s13578-023-01132-1.

110. Zohar, O., Reiter, Y., Bennink, J.R., Lev, A., Cavallaro, S., Paratore, S., Pick, C.G., Brooker, G., and Yewdell, J.W. (2008). Cutting edge: MHC class I-Ly49 interaction regulates neuronal function. J Immunol 180, 6447–6451. 10.4049/jimmunol.180.10.6447.

111. Adelson, J.D., Sapp, R.W., Brott, B.K., Lee, H., Miyamichi, K., Luo, L., Cheng, S., Djurisic, M., and Shatz, C.J. (2016). Developmental Sculpting of Intracortical Circuits by MHC Class I H2-Db and H2-Kb. Cereb Cortex 26, 1453–1463. 10.1093/cercor/bhu243.

112. Estes, M.L., and McAllister, A.K. (2015). Immune mediators in the brain and peripheral tissues in autism spectrum disorder. Nat Rev Neurosci 16, 469–486. 10.1038/nrn3978.

113. Estes, M.L., and McAllister, A.K. (2016). Maternal immune activation: Implications for neuropsychiatric disorders. Science 353, 772–777. 10.1126/science.aag3194.

114. McAllister, A.K. (2017). Immune Contributions to Cause and Effect in Autism Spectrum Disorder. Biol Psychiatry 81, 380–382. 10.1016/j.biopsych.2016.12.024.

115. Zalocusky, K.A., Najm, R., Taubes, A.L., Hao, Y., Yoon, S.Y., Koutsodendris, N., Nelson, M.R., Rao, A., Bennett, D.A., Bant, J., et al. (2021). Neuronal ApoE upregulates MHC-I expression to drive selective neurodegeneration in Alzheimer’s disease. Nature neuroscience 24, 786–798. 10.1038/s41593-021-00851-3.

116. Tan, C.X., Burrus Lane, C.J., and Eroglu, C. (2021). Role of astrocytes in synapse formation and maturation. Curr Top Dev Biol 142, 371–407. 10.1016/bs.ctdb.2020.12.010.

